# A Unified Approach to Investigating 4 dpf Zebrafish Larval Behaviour through a Standardised Light/Dark Assay

**DOI:** 10.1101/2023.11.23.568441

**Authors:** Courtney Hillman, James Kearn, Matthew O. Parker

## Abstract

Zebrafish have emerged as a dynamic research model in the domains of neuropsychopharmacology, biological psychiatry and behaviour. Working with larvae ≤ 4 days post-fertilisation (dpf) offers an avenue for high-throughput investigation whilst aligning with the 3Rs principles of animal research. The light/dark assay, which is the most used behavioural assay for larval neuropharmacology research, lacks experimental reliability and standardisation. This study aimed to formulate a robust, reproducible and standardised light/dark behavioural assay using 4 dpf zebrafish larvae. Considerable between-batch and inter-individual variability was found, which we rectified with a normalisation approach to ensure a reliable foundation for analysis. We then identified that 5-minute light/dark transition periods are optimal for locomotor activity. We also found that a 30-minute acclimation in the light was found to produce significantly increased dark phase larval locomotion. Next, we confirmed the pharmacological predictivity of the standardised assay using ethanol which, as predicted, caused hyperlocomotion at low concentrations and hypolocomotion at high concentrations. Finally, the assay was validated by assessing the behavioural phenotype of hyperactive transgenic (*adgrl3.1*^-/-^) larvae, which was rescued with psychostimulant medications. Our standardised assay not only provides a clear experimental and analytical framework to work with 4 dpf larvae, but also facilitates between-laboratory collaboration using our normalisation approach.

## 1. Introduction

Zebrafish (*Danio rerio,* Hamilton 1822) larvae are an increasingly popular research model with a breadth of application and tractability for neuropsychopharmacology (Cassar et al., 2020). Due to their rapid breeding and small size, working with larvae provides potential for high-throughput approaches, with the added benefit that larvae up to 5 days post-fertilisation (dpf) are exempt from protection by national legislation (e.g., the (Animals (Scientific Procedures) Act 1986) 1986) in the United Kingdom (UK) and the European Union (EU) Directive 2010/63/EU of the European Parliament. Hence, 4 dpf larval research has significant interest for meeting the criteria of the 3Rs (replacement, refinement and reduction) of *in vivo* animal research (Crilly et al., 2018). Indeed, as a first-tier model, 4 dpf zebrafish larvae can reduce the number of protected animals required at later testing stages as well as minimising the suffering of experimental animals when testing toxic substances. Therefore, from a 3Rs perspective, zebrafish larvae are a unique and appealing research model.

The light/dark assay remains the most used for studying the behaviour of zebrafish larvae owing to its cost and time-effectiveness, as well as its high face and pharmacological predictivity (Haigis et al., 2022). The protocol manipulates the innate tendency for larvae to display hyperactivity following a sudden change from light to dark conditions (Bachour et al., 2020). Some have suggested that this locomotor response represents a stress or alarm response, and it has been used for modelling anxiety-like phenotypes in larvae (Rock et al., 2022; Steenbergen et al., 2012). Its popularity stems from its ease of manipulation for toxicity and drug analysis and the robust behavioural response observed (Chatzimitakos et al., 2022; Haigis et al., 2022; Kirla et al., 2021; Licitra et al., 2021; Nixon et al., 2021; Pesavento et al., 2022; Sales Cadena et al., 2021; Zaig et al., 2021). As seen in Table 1, there is, however, a lack of consistency between light/dark assay experimental protocols in 4 dpf larvae. The lack of experimental consistency can impact the behavioural responses observed and hence the reproducibility of results between laboratories and researchers (Pesavento et al., 2022). This lack of standardised testing and variability in protocols has been highlighted previously by Legradi (2015). However, no standardised or reproducible assay has been published to date.

**Table 1.**
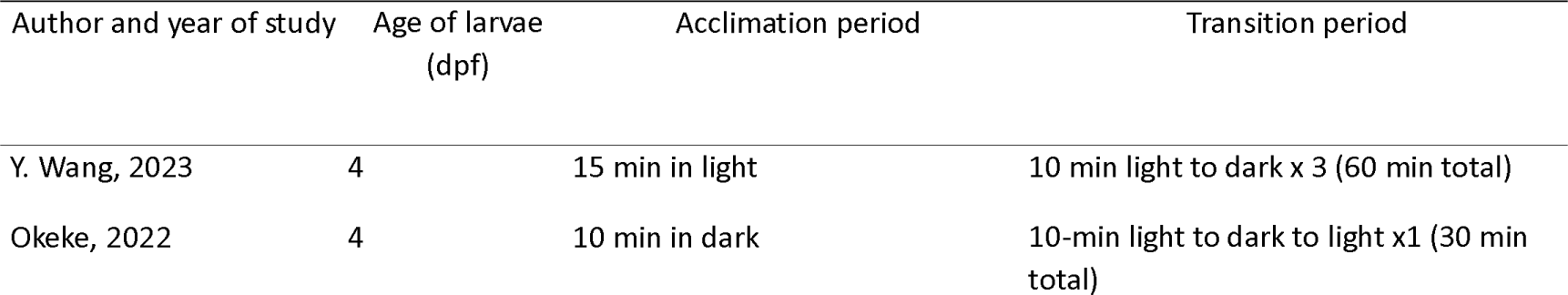
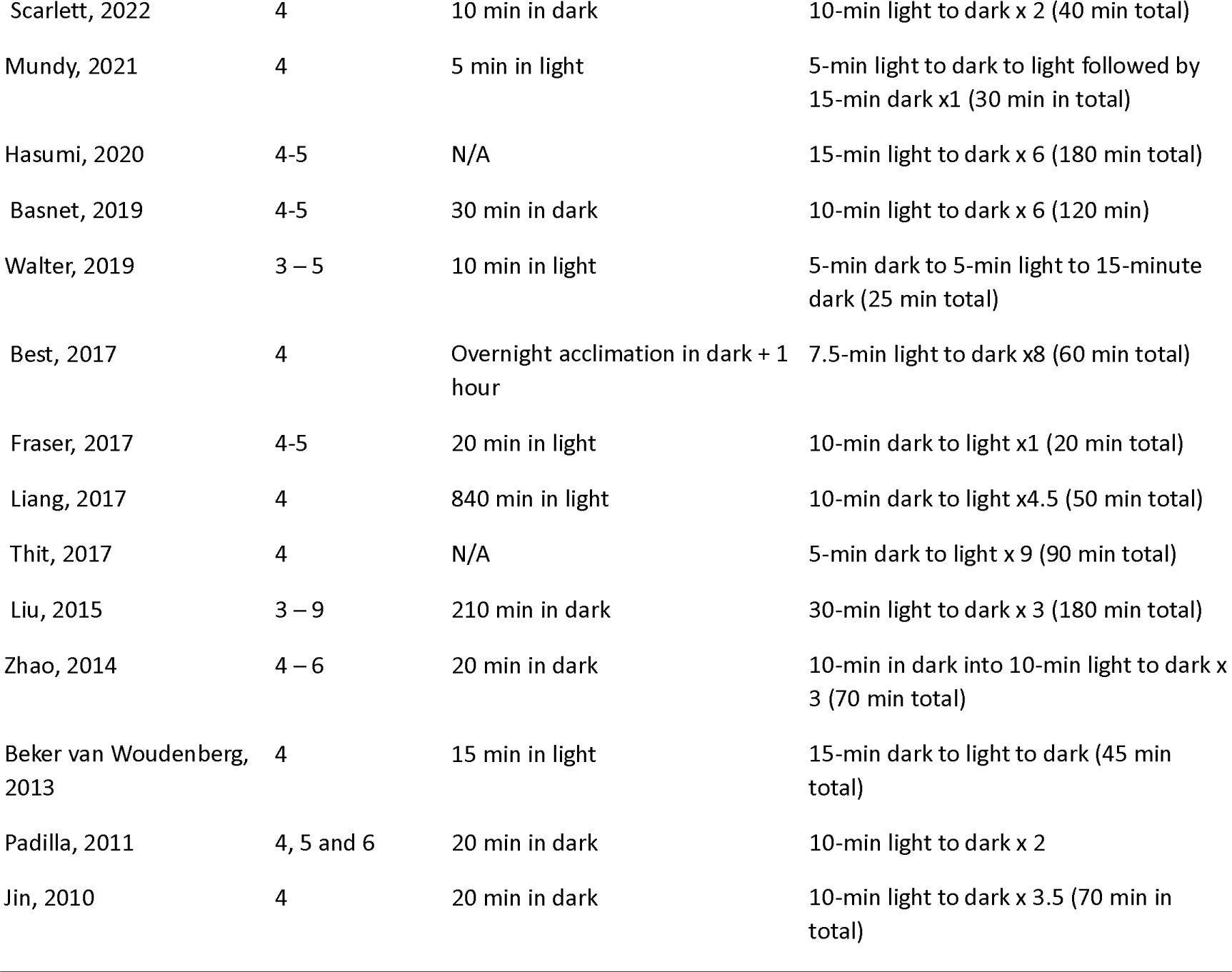
List of previous publications using 4 dpf zebrafish larvae and the different light dark parameters used. dpf: days post-fertilisation, min: minutes.

Here, we aimed to develop a standardised robust and reproducible light/dark behavioural assay using 4 dpf zebrafish larvae and to validate this assay using pharmacological and genetic manipulations. After assessing different parameters to determine the optimal experimental conditions, we validated the assay pharmacologically in two-ways. Wild-type larvae were exposed to ethanol and genetically modified (*adgrl3.1* knock-out *(adgrl3.1*^-/-^)) larvae were used to study behavioural interventions of two Attention-Deficit Hyperactivity Disorder (ADHD) drugs. Ethanol was selected due to its well-known ability to induce low concentration hyperactivity and high concentration hypoactivity, which we postulated would impact the dark phase responses (McLellan, 2017). Furthermore, it is widely available, is globally abused, and is commonly used in larval light/dark assay research (Davis et al., 2021; Irons et al., 2010; MacPhail et al., 2009). The *adgrl3.1*^-/-^ mutant line demonstrates phenotypes typical of ADHD and therefore we hypothesised that the larvae would display hyperactivity in the dark phase (Sveinsdóttir et al., 2023) which would be rescued/reversed with methylphenidate (MPH), a standard stimulant drug used to treat ADHD. We also assessed atomoxetine (ATO), another common ADHD drug known to induce sedation in patients, which we postulated would result in hypoactivity (Bastiaens, 2007). In addition to the validation, we also aimed to provide a consolidated method of analysis and interpretation of the behavioural data, including radar charts and heat maps of different swim patterns, which will be of use to the zebrafish research community.

## 2. Materials and Methods

### 2.1. Animal husbandry

AB wild-type (WT) zebrafish were bred in-house from the University of Portsmouth colony. Breeding adults in the colony were housed in ∼50:50 male:female groups of 8-10 per 2.8 L tank, or up to 20 per 6 L tank, on a re-circulating system (Aquaneering, Fairfield Controlec Ltd., Grimsby, UK). Sample sizes (n = 16) were calculated based on power analyses from effect sizes observed in our pilot studies (*see section 2.8. Power Analysis*). *adgrl3.1*^-/-^ larvae were used for the genetic phenotype experiment. The *adgrl3.1*^-/-^ line was created by CRISPR-Cas9 genome engineering as previously described (Sveinsdóttir et al., 2023) and maintained via homozygous in-cross at the University of Portsmouth fish facility (with matched-age WT [AB] controls). Room and tank water temperatures were maintained at 25-27°C, pH 8.4 (±0.4), on a 14:10-hour light/dark cycle with the lights switched on at 08:30. Fish were fed on ZM-100 fry food (Zebrafish Management Ltd., Winchester, UK) from 5 dpf until adulthood when they were moved onto a diet of flake food and live brine shrimp 3 times/day (once/day on weekends). Larvae were bred for experimentation either by adding marbles to the base of the tank overnight (to induce spawning) or by pair breeding. The larvae used in these experiments originated from over 10 different batches from different breeders. Larvae were reared in a translucent incubator (28.5°C) in the fish facility until the day of testing, where they were moved into our behavioural testing room and placed in a small countertop incubator at 28°C. On completion of behavioural testing, larvae were killed using Aqua-Sed (Aqua-Sed^TM^, Vetark, Winchester, UK) in accordance with manufacturer guidelines. All experiments were approved by the University of Portsmouth Animal Welfare and Ethical Review Board. Genetically altered fish were used under license from the UK Home Office (Animals (Scientific Procedures) Act, 1986, Licence number PP8708123).

### 2.2. Behavioural apparatus

All behavioural testing was carried out between 09:00 and 18:00 (Mon-Fri) in Zantiks MWP units (Zantiks Ltd, Cambridge, UK). Previous literature determined that 48-well plates are the optimal format for consistent and robust behavioural responses and high-throughput analysis (Widrick et al., 2023). However, our preliminary studies suggest any well-plate size is suitable *(see SD 1).* Therefore, with respect to previous literature and both throughput and time-efficiency, all behavioural testing took place using 48 well plates. Filming was carried out directly from above the plates, which allowed live monitoring within the behavioural system.

### 2.3. Standardisation of Light/Dark Assay

A breakdown of the experimental procedure is seen in Figure 1. At 08:30 the larvae Petri dishes were transferred into an incubator in our behavioural testing room at 28°c to allow the fish to acclimate to the new environment for 30 minutes, prior to experimentation. 225 µL of Petri dish water and a randomly selected larva were transferred into each well of a 48-well plate using trimmed pipette tips (∼ 2 mm removed from the tip end of a 1000 µL pipette tip). The larvae were selected from different Petri dishes at random for each experiment. The well plate was transferred to the Zantiks MWP unit, and the overhead lights in the unit were switched on for 30-minutes for acclimation (Fig. 1A). Following acclimation, the light/dark tracking script was run for a 30-minute baseline recording. This comprised three sets of five-minute light phase (white light, 350 lux) to dark phase transitions (Fig. 1B). The MWP unit was temperature controlled throughout experimentation at 28.02°c ± 0.07.

**Figure 1.**
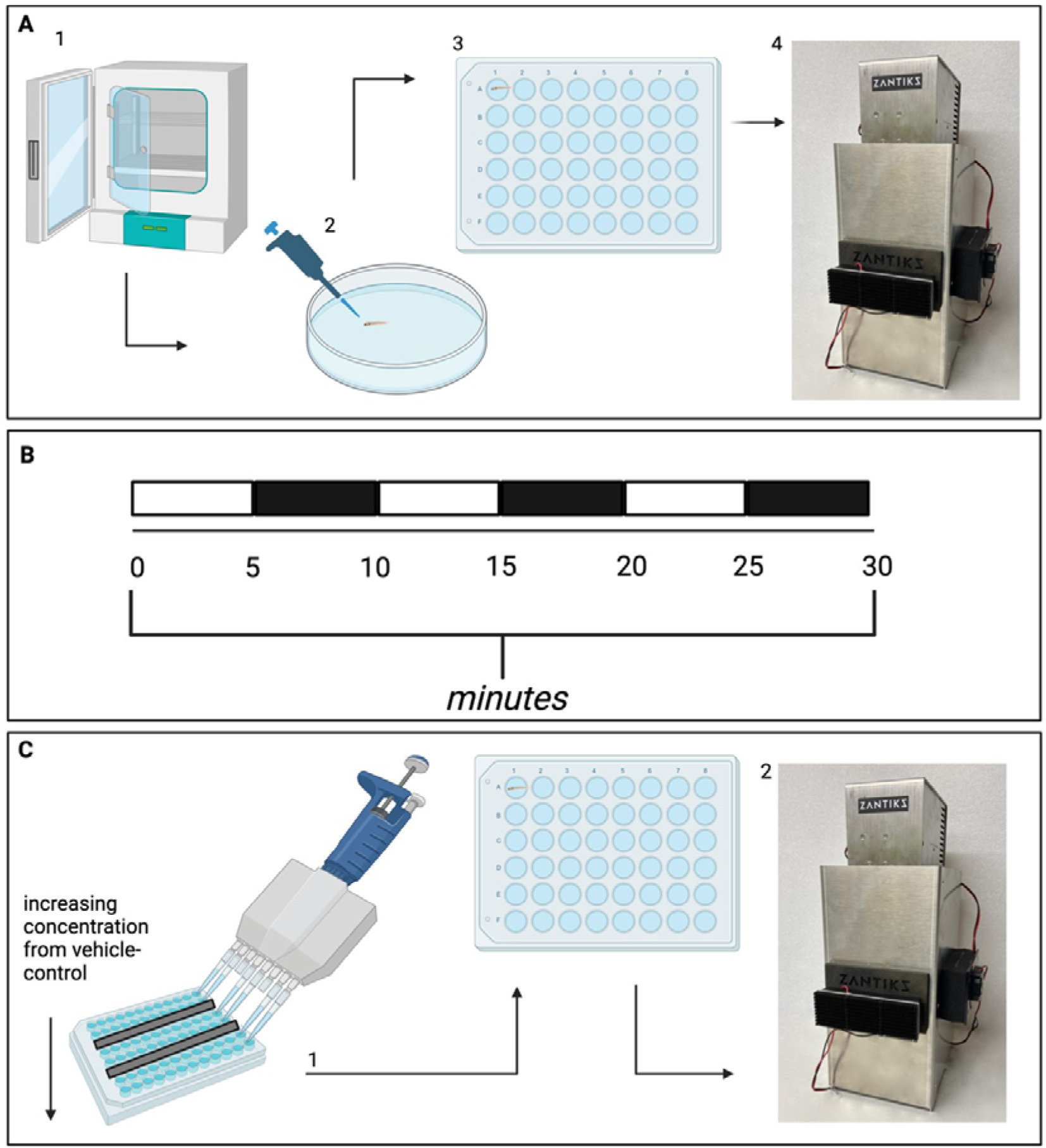
The experimental protocol for the standardised light/dark assay including drug exposure procedures. (A) the fish are transferred from the incubator to a 48 well plate and placed in the behavioural unit with the lights on for 30-minutes of acclimation prior to the recording. (B) Breakdown of the L/D (light/dark) recording which incorporates 5-minute light phase to dark phase transitions for 30-minutes. (C) Overview of drug exposure procedure from a 96-well dosing plate to the fish. Behaviour was then monitored using a Zantiks MWP unit.

### 2.4. The Light/Dark Assay for Chemical Exposures

The drug exposure protocol incorporated the standardised protocol (*Section 2.3 of this paper*). Following baseline recording, the plate was removed from the MWP, and a multi-channel pipette was used to administer 25 µL of different ethanol concentrations *(CAS number: 64-17-5, tested at 0.5 – 4.0 %)* (Merck Life Sciences UK Ltd., Gillingham, UK) from a 96-well plate to achieve the desired final exposure concentration (96-well plate contained 5-40% concentrations) (Fig.1C). The 48-well plate was returned to the unit and the larvae were tracked during the light/dark alternations (Fig 1B).

### 2.5. The Light/Dark Assay for Genomic Phenotype Evaluation

Assays with adgrl3.1^-/-^ larvae incorporated the standardised protocol and both WT and adgrl3.1^-/-^ larvae were used (*Section 2.3 of this paper*). Following the baseline recording, the plate was removed, and a multi-channel pipette was used to administer 0.05 – 20 μM MPH *(CAS number: 298-59-9)* (Merck Life Sciences UK Ltd., Gillingham, UK) *and* 2.93 - 10.2 μM ATO *(CAS number: 82248-59-7)* (Merck Life Sciences UK Ltd., Gillingham, UK) (Fig. 1C). Both MPH and ATO were initially diluted in 100% dimethyl sulfoxide (DMSO) for a final concentration of 0.5%, which is a safe concentration in larvae (Hoyberghs et al., 2021). The 48-well plate was returned to the unit and the larvae were subjected to the light/dark protocol (Fig 1B).

### 2.6. Optimisation of the Light/Dark Assay

Alterations in a range of parameters were investigated as part of the standardisation of the light/dark assay. This included assessing previously published protocols (Acevedo-Canabal et al., 2019; Bachour et al., 2020; Haigis et al., 2022), for determining the effect of altering the length of light/dark transition periods. The length of acclimation period and lighting conditions were assessed as well as the effect of the time of day upon behavioural responses.

### 2.7. Statistics

The data files were imported into a custom R processing script (R Core Team, 2021) where they were normalised to baseline to overcome larval batch variability. Normalisation was carried out by determining the percentage of movement compared to the overall baseline mean response of all the larvae. Outliers were identified and eliminated if they were greater than the absolute deviation around the median, a method commonly used in neuropsyhcopharmacological research for outlier identification and removal (Leys et al., 2013). All graphs and analyses were produced and performed using the R script. Data was assessed for distribution using the Shapiro-Wilk test and Levine’s test for homogeneity. Statistically significant differences were assessed using a one-way analysis of variance (ANOVA) and for chemical/ strain analyses, linear mixed models were implemented with movement as the dependent variable, concentration or strain of fish, light phase and time as independent variables and fish ID as the random effect. A second linear mixed model analysis was run on the dark phase data only with movement as the dependent variable, concentration or strain of fish and time as independent variables and fish ID as the random effect. Likelihood ratio tests (LRT) were performed on all models (i.e., the fitted models compared to null models which included only random effects) to test the null hypotheses that added parameters (independent variables) did not have a significant effect on the outcomes. Post-hoc tests incorporated into the analysis included Tukey’s multiple comparison for comparing dark phase responses and linear regressions analysing dark phase movement with time as the independent variable. The alpha levels are P*** < 0.001, P*\*\** < 0.01, P*\** < 0.05.

Radar charts and heat maps were produced by adapting an approach described by Haigis (2022) and Steele (2018). Briefly, within the custom R processing script, the percentage of baseline data was analysed as either darting (>2000 %/s), steady swimming (690 – 2000 %/s), slow swimming (0.1 - 690 %/s) or zero (0 %/s) movements, the values of which were adapted from Haigis (2022), Steele (2018) and Scarlett (2022) according to relative movement in our experiments. Each observation had an addition of one to overcome the limitations of percentage determination using a zero response. The endpoints evaluated were total movements, darting swimming movements, steady swimming movements, slow swimming movements and the absence of movements (zero movements). The mean response for each of these observations was calculated for controls. The percentage of movement compared to vehicle-controls was calculated for each drug concentration and a base 10 logarithmic conversion was performed.

### 2.8. Power Analysis

Observed power and where appropriate, sample size, was calculated for all tested concentrations and strains of fish. When observed power was below 0.8 (80%), a sample size calculation was performed. If the observed power was ≥ 0.8, then the sample size was n = 16. Figure 2 provides an overview of this. All power analysis results, code, data and comprehensive statistical results can be found at the OSF linked to this paper: https://osf.io/k7y2c/?view_only=61e23ea4bb2d452dab76646931e3f332.

**Figure. 2.**
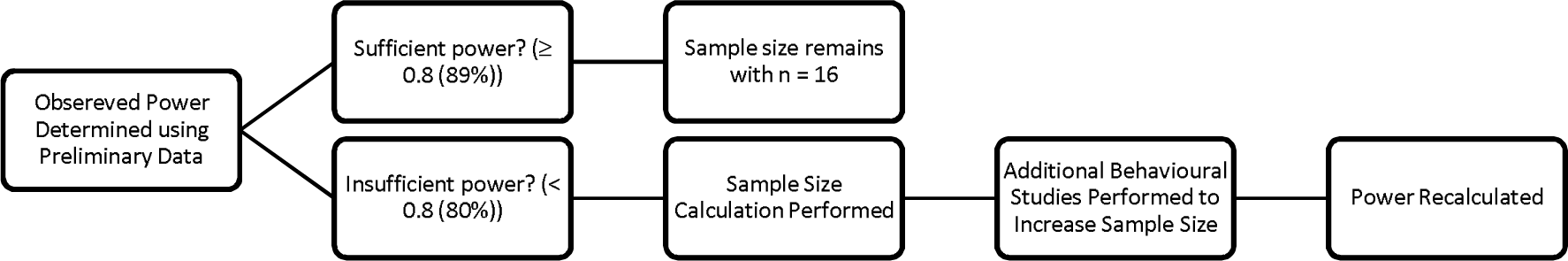
Flow chart depicting the power analysis and sample size determination procedure for larval zebrafish drug exposures.

All figures were created with Biorender.com and data is represented as mean ± SEM. The minute analysis graphs have data points which represent averaged locomotor responses of the fish. Comparatively, the bar charts have data points which represent individual fish responses.

## 3. Results

### 3.1. Five-minute light/dark protocol produces a robust response in 4 dpf zebrafish larvae allowing for high throughout behavioural analysis

Previous publications have lacked consistency with respect to the length and number of light/dark transition periods for assays using 4 dpf zebrafish larvae (Table 1). Therefore, we assessed the optimal length of light to dark transition periods by replicating the protocols employed by Acevedo-Canabal (2019), Bachour (2020) and Haigis (2022), as well as assessing our own protocol. There were reproducible and robust dark phase locomotor responses with all transition periods. In addition, the five-minute (one-way ANOVA: F_(2, 41)_ = 0.656, P = 0.524) and ten-minute transition (one-way ANOVA: F_(3, 60)_ = 0.21, P = 0.889) protocols, had no significant change in locomotion across successive dark phases (Fig. 3D and 3F). In both the two-minute (one-way ANOVA: F_(4, 75)_ = 3.654, P = 0.008) and fifteen-minute transition (one-way ANOVA: F_(5, 86)_ = 2.731, P = 0.024) protocols, however, we observed a significant reduction across successive dark phases (Fig. 3B and 3H). With the two-minute transition period, locomotion in dark phases four and five is significantly less than dark phase one (Fig. 3B). Similarly, with the fifteen-minute transition period, dark phase six is significantly less than dark phase one (Fig. 3H).

**Figure 3.**
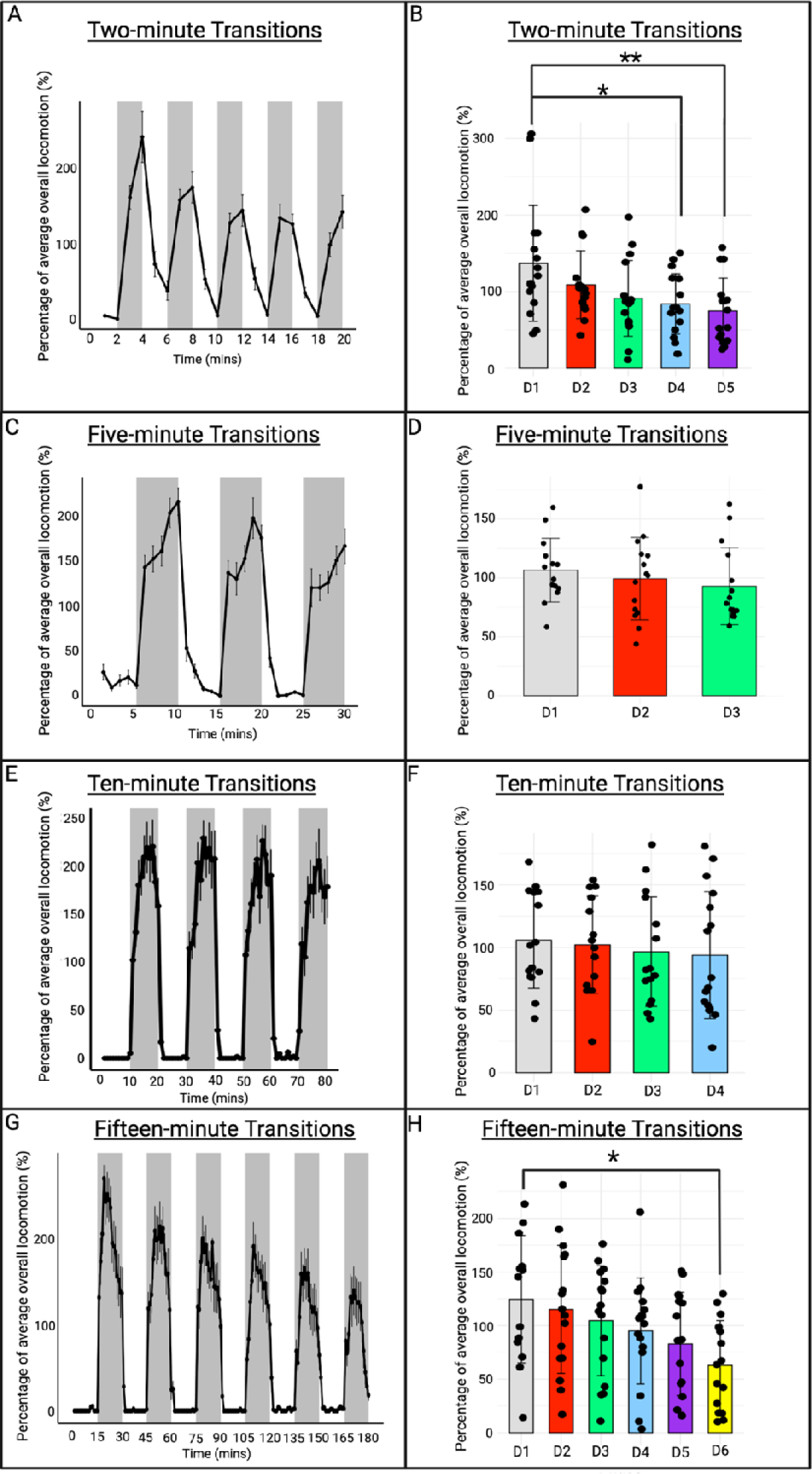
Effect of varying light and dark phase length on larval behavioural response. Light/dark locomotion displayed as a percentage of baseline response are represented in minutes for (A) two-minute transitions (C) five-minute transitions (E) ten-minute transitions and (G) fifteen-minute transitions. The mean dark phase responses are numbered as D1-D6 for (B) two-minute transitions, (D) five-minute transitions, (F) ten-minute transitions and (H) fifteen-minute transitions.

We next looked in detail at locomotion within the dark phases, and calculated the locomotion within the first minute, half-way through, and during the final minute of the dark phase (Fig. 4) (n.b., the two-minute period was only sufficient for a single dark phase analysis). There was a significantly reduced initial dark phase locomotion during the fifteen-minute protocol compared to the two-minute transition period (one-way ANOVA: F_(3, 60)_ = 4.002, = 0.0115) (Fig. 4A). No significant difference in locomotion is seen half-way through the dark phase regardless of length of transition period (one-way ANOVA: F_(2, 44)_ = 2.543, = 0.0901) (Fig. 4B). However, by the end of the recording period, the two-minute, five-minute and ten-minute locomotion’s are significantly greater than the fifteen-minute protocol (one-way ANOVA: F_(3, 60)_ = 7.021, = 0.0004) (Fig. 4C).

**Figure 4.**
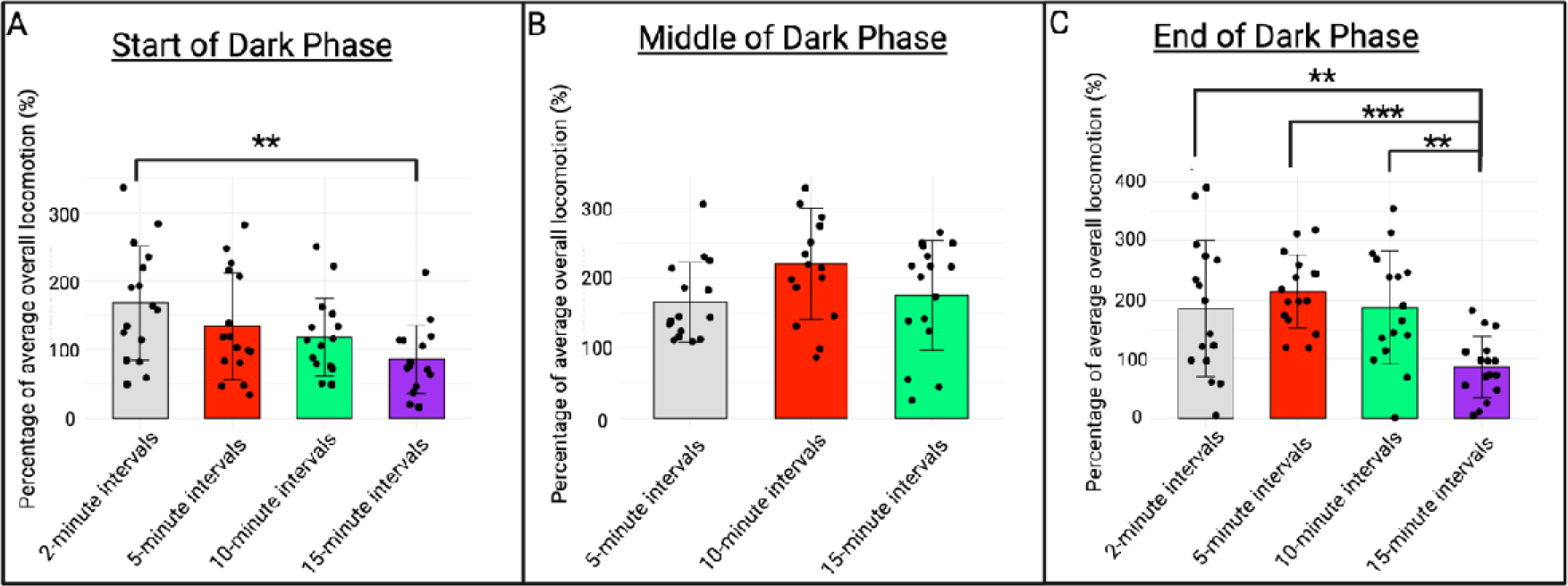
Time-dependent effects of varying light and dark phase length on larval behavioural responses. Locomotion displayed as a percentage of baseline response represented as mean dark phase responses for (A) the first minute of each dark phase (B) the median minute of each dark phase and (C) the final minute of each dark phase for the two-minute, five-minute, ten-minute and fifteen-minute transition protocols.

In view of the sustained increased locomotion throughout the recording period with the five-minute transition protocol, and relatively higher throughput potential, we proceeded with further testing using the five-minute transition protocol.

### 3.2. The acclimation period affects locomotor behavioural responses in the larval light/dark assay

The importance of the length and lighting conditions of the acclimation period has been previously reported (Makaras et al., 2021). Larval light/dark research lacks standardisation of the length and lighting conditions of the acclimation period (Table 1). The effects of the length and lighting conditions of acclimation within the behavioural monitoring unit on light/dark assay locomotor responses were addressed by comparing no acclimation, five-minute, ten-minute, 20-minute or 30-minute acclimation periods (Fig. 5), as well as varying the lighting conditions (Fig. 6). Our findings in Figure 5 show a significant effect of acclimation period length on locomotor responses (one-way ANOVA: F_(4, 75)_ = 7.249, < 0.0001). 30-minute acclimation produced significantly greater dark phase locomotion compared to the other tested acclimation periods (Fig. 5F). Therefore, we chose a 30-minute acclimation for all further experimentation due to the increased dark phase responsiveness.

**Figure 5.**
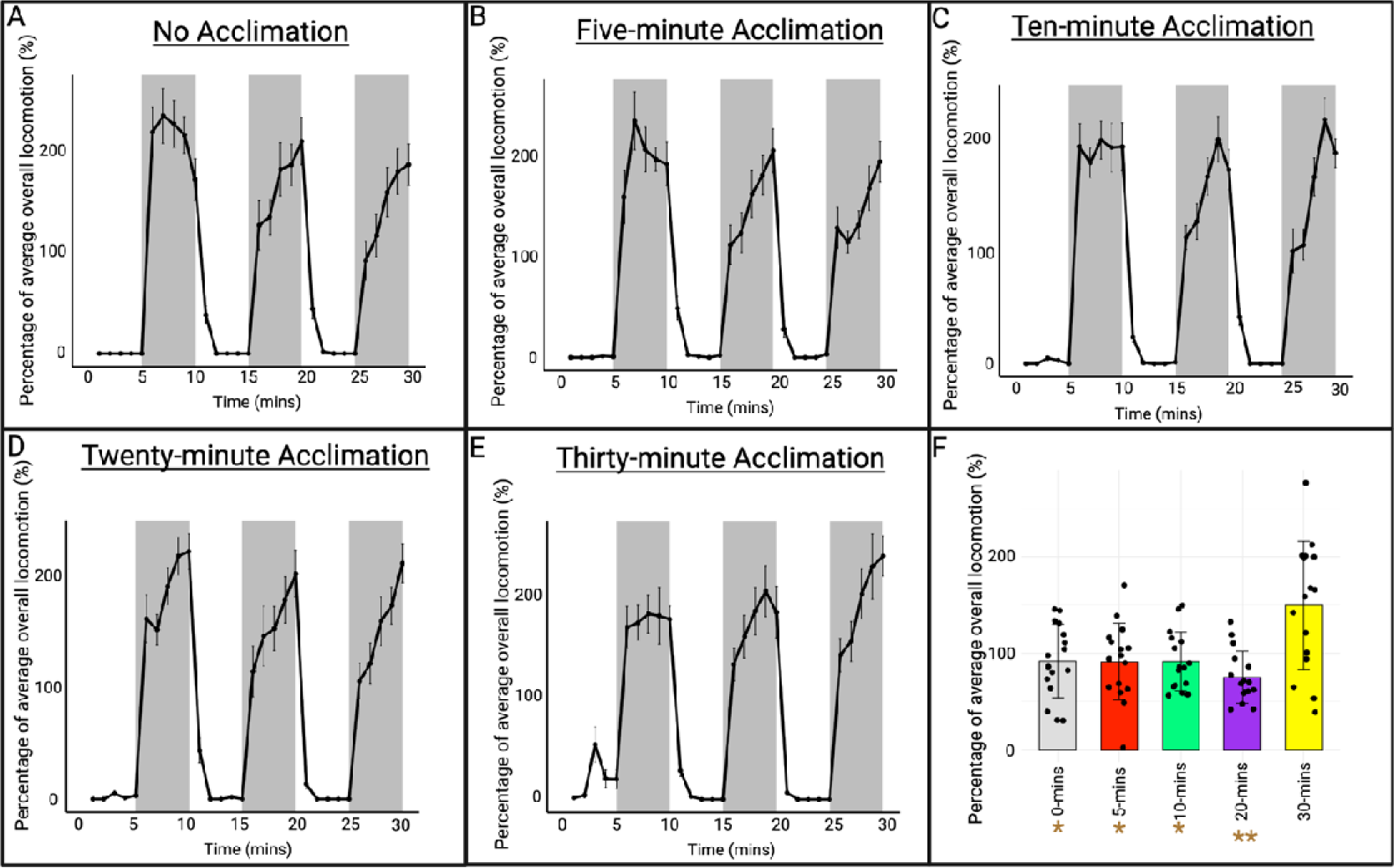
The light/dark response is unaffected by acclimation period length. The light/dark mean locomotor responses represented in minutes for (A) no acclimation, (B) five-minute acclimation in the light, (C) ten-minute acclimation in the light, (D) 20-minute acclimation in the light and (E) 30-minute acclimation in the light. (F) represents the mean dark phase locomotor responses for each acclimation period length.

**Figure 6.**
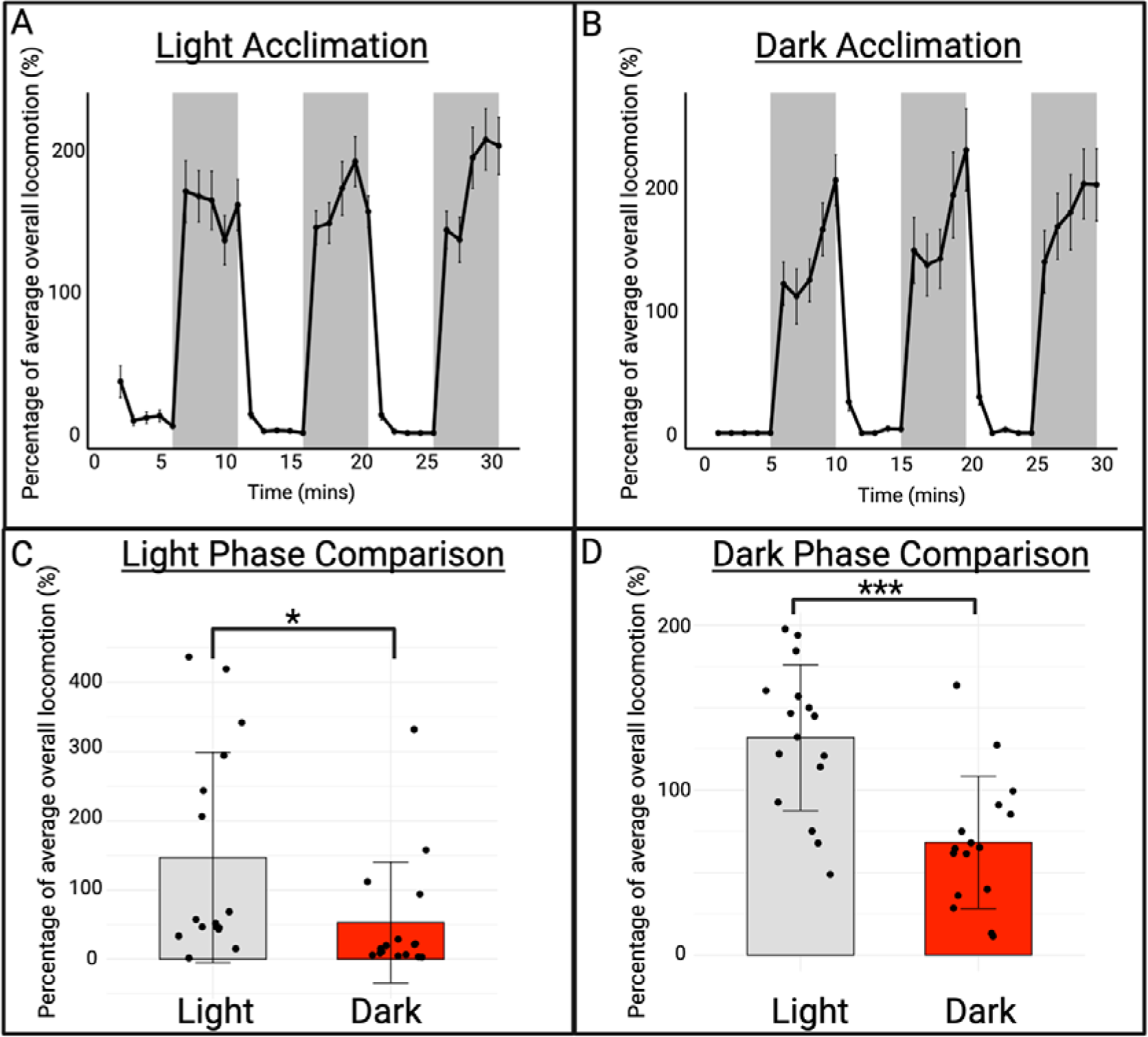
Lighting conditions in the acclimation period do not affect the subsequent light/dark assay behavioural response. Larvae were acclimated in the light (A) or dark (B) for 30 minutes and then underwent the standardised light/dark assay (section 2.3 of this paper).

Varying the lighting conditions (light or dark) during the acclimation period revealed a significant effect on light (one-way ANOVA: F_(1, 30)_ = 4.611, = 0.04) and dark phase (one-way ANOVA: F_(1, 30)_ = 18.09, < 0.0001) locomotion (Fig. 6). Therefore, due to the increased overall locomotion, light acclimation was selected for further experimentation.

### 3.3. The light/dark assay is suitable for assessing the effects of chemicals on larval behaviour

We next validated the assay in the context of behavioural pharmacology by examining larval behavioural responses to different concentrations of ethanol. Ethanol was selected due to its widespread availability and common use in larval light/dark assay research (Davis et al., 2021; Irons et al., 2010; MacPhail et al., 2009). We also sought to identify meaningful and effective interpretations of the light/dark assay data (Fig. 7). By applying these straightforward analyses, distinctive concentration-dependent altered locomotor responses in both light and dark phases were revealed.

**Figure 7.**
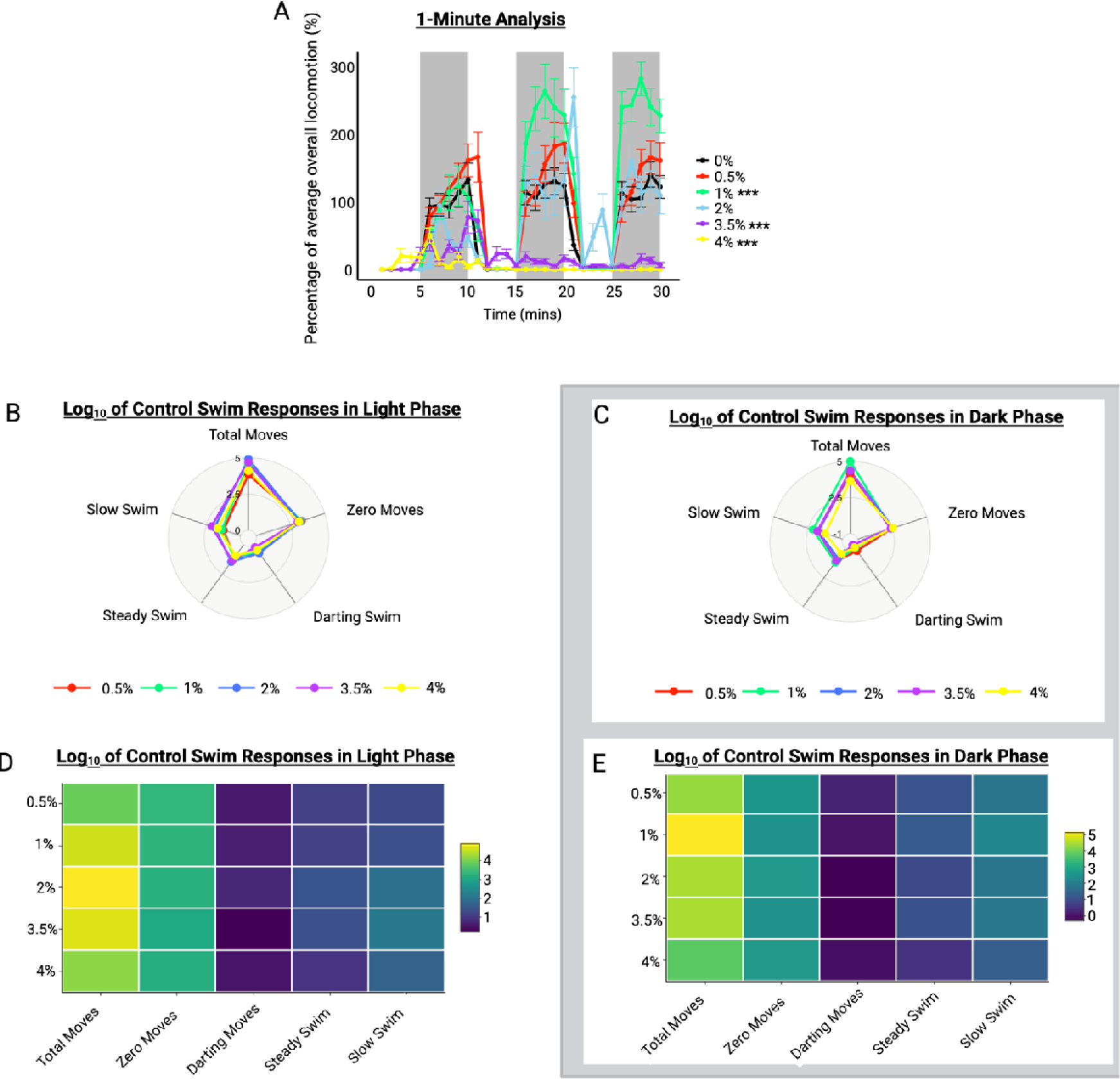
Concentration-dependent effects of ethanol on larval behaviour in the light/dark assay. The minute analysis (A) is represented as mean percentage of baseline movement over the 30-minute recording period. The heat map (B) and radar chart (D) show light phase responses and (C) and (E) show dark phase responses.

Figure 7A shows that ethanol exposure has a significant effect upon both light and dark phase locomotion (linear mixed model analysis (lights effect): β = 81.58, SE = 3.30, t = 24.76). To compare this model to the null model, an LRT was conducted. The LRT showed a significant difference between the fitted model and the null model, 2(1) = 543.11, < 0.0001. Specifically, low concentrations of ethanol (1.0%) increase locomotion in the dark phase compared to control larvae. 3.5% and 4.0% exposed larvae had decreased locomotion in the dark phase compared to controls (Fig. 7A). The fish exposed to the 2.0% ethanol appeared to increase locomotion in the light phase compared to control larvae with relatively unchanged dark phase locomotion. We hypothesised that this altered locomotion was reduced responsiveness to the light/dark stimuli (Fig. 8). However, further analysis disregarded our hypothesis with 2.0% ethanol exposed larvae having no significant difference in light phase locomotion compared to controls following a student’s t-test analysis (Paired t-test, t(15) = -1.3924, = 0.1847).

**Figure 8.**
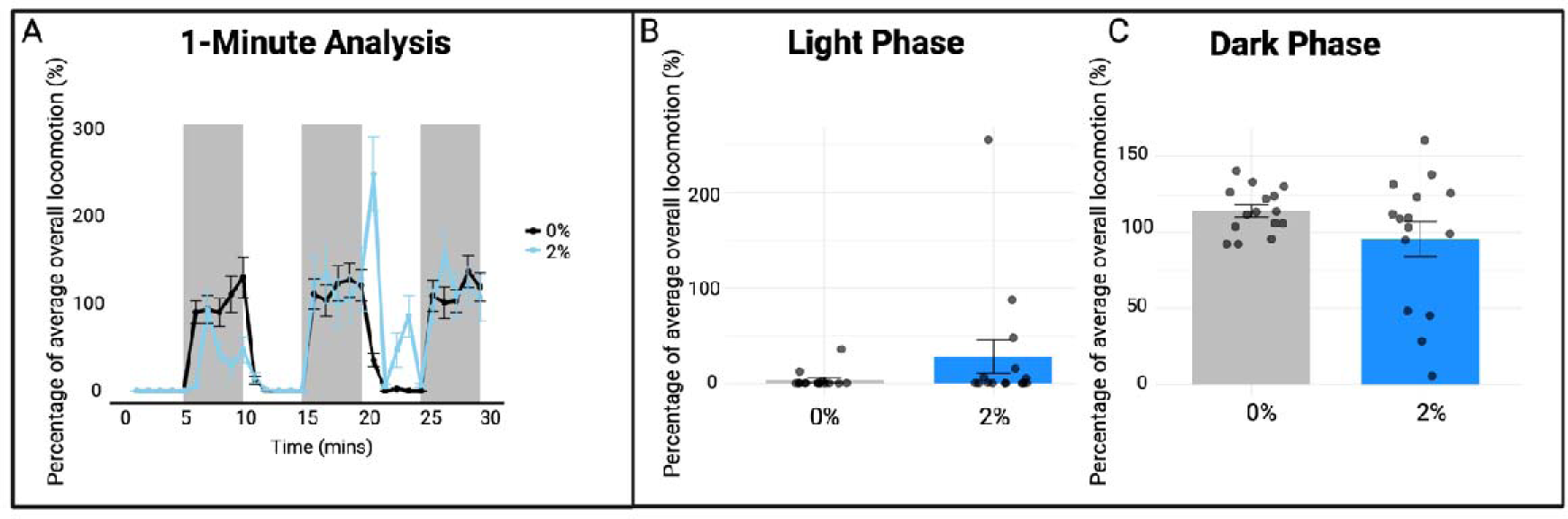
Increased light phase responsiveness with exposure to 2.0% ethanol in zebrafish larvae. The minute analysis (A) and light vs dark average (B and C) are represented as mean percentage of baseline movement over the 30-minute recording period.

To gain more intricate concentration-dependent behavioural responses, we next furthered our analysis and interpretation of the data by adapting heat maps and radar charts (Haigis (2022) and Steele (2018)) to the larval light/dark analysis. The type of swim movement that is increased following addition of low concentrations of ethanol is ‘total movements’ (one-way ANOVA: F_(5, 90)_ = 0.856, = 0.514) (Fig. 7C and E). 4.0%-exposed larvae had equivalent darting responses to control larvae; however, they had reduced total and steady movements (Fig. 7D and E). Furthermore, in the light phase all concentrations had reduced total movement compared to control larvae (one-way ANOVA: F_(5,90)_ = 1.393, = 0.235) (Fig. 7B and D). 2.0%-exposed larvae had increased darting responses compared to controls and 4.0%-exposed fish had increased slow swim responses compared to the other concentrations. These specific swim observations we have highlighted here are not seen on the line graph and bar chart. Therefore, the radars and heat maps facilitate interpretation of the data and provide insight to more advanced swim responses which cannot be achieved with basic movement analyses.

Next, using the same data, we examined time-dependent effects of ethanol exposures for the first 10-minutes, 10-20 minutes and 20-30 minutes post-exposure (Fig. 9). By the first dark phase, there is a significant reduction in locomotion with 4% ethanol compared to controls (one-way ANOVA: F_(5, 87)_ = 5.55, < 0.0001) (Fig. 9B). The significant increase in locomotion with 1.0% ethanol is not observed until the second dark phase (one-way ANOVA: F_(5, 84)_ = 13.02, < 0.0001) (Fig. 9D). 4.0% ethanol exposed larvae had reduced dark phase locomotion as well (Fig. 9D). By the final 10-minutes, the 2.0% exposed larvae showed increased light phase locomotion (one-way ANOVA: F_(5, 80)_ = 14.45, < 0.0001) (Fig. 9E). This demonstrates the potential of the assay to examine behavioural expressions of pharmacokinetics/dynamics and highlights the importance of evaluating time-dependence in addition to overall recording responses over the full 30-minutes.

**Figure 9.**
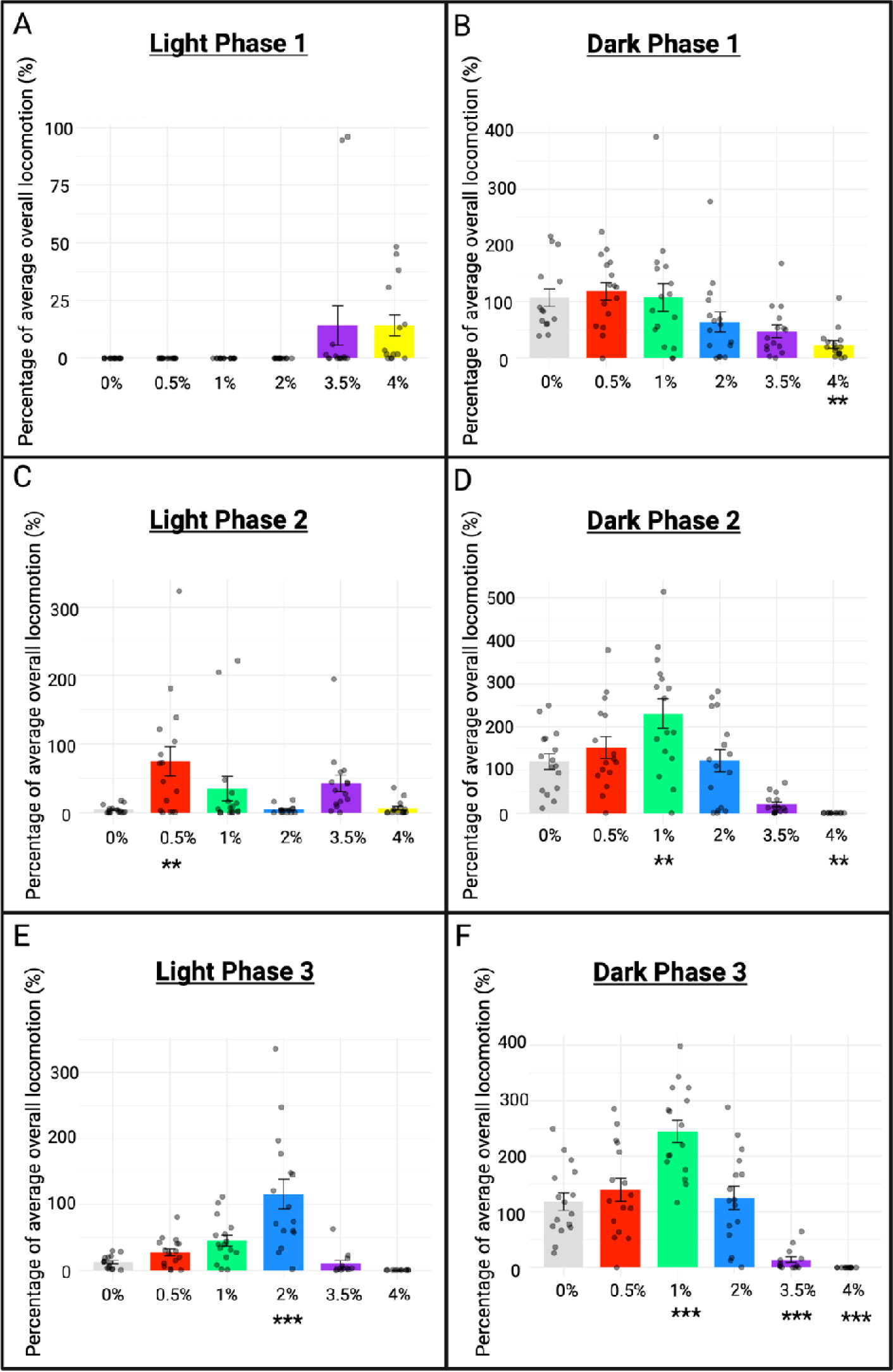
Time-dependent effects of ethanol exposure on larval light/dark behaviour. Mean locomotor responses represented as a percentage of baseline movement for (A and B) light and dark phase one (0-10-minutes post-exposure), (C and D) light and dark phase two (10-20-minutes post-exposure) and (E and F) light and dark phase three (20-30-minutes post-exposure).

### 3.4. The light/dark assay is suitable for assessing behavioural phenotypes of transgenic zebrafish lines

We next validated the assay by examining the phenotypes of *adgrl3.1*^-/-^ larvae, which have previously been shown to express hyperactivity (Sveinsdóttir et al., 2023). In previous work, the noradrenergic reuptake inhibitor, MPH, a therapeutic drug that reduces hyperactivity in children with ADHD, has been shown to rescue the hyperactive phenotype. We examined the effects of this drug on the light/dark assay responses at concentrations determined from previous publications (Lange et al., 2012).

Our findings in figure 10 show a significant increase in dark phase locomotion in the *adgrl3.1*^-/-^ larvae compared to WT larvae, which is partially rescued following addition of 5 and 20 μM MPH (linear mixed model analysis (lights effect): *β* = 140.24 (SE = 2.6, *t* = 53.97) (Fig. 10A). The LRT showed a significant difference between the fitted model and the null model, X2 (7) = 2085.4, P < 0.0001. No significant changes were observed for the WT larvae treated with 0.05 μM and 5 μM MPH compared to untreated WT fish in either light condition, but 20 μM MPH significantly reduced dark phase responses (Fig. 10A).

**Figure 10.**
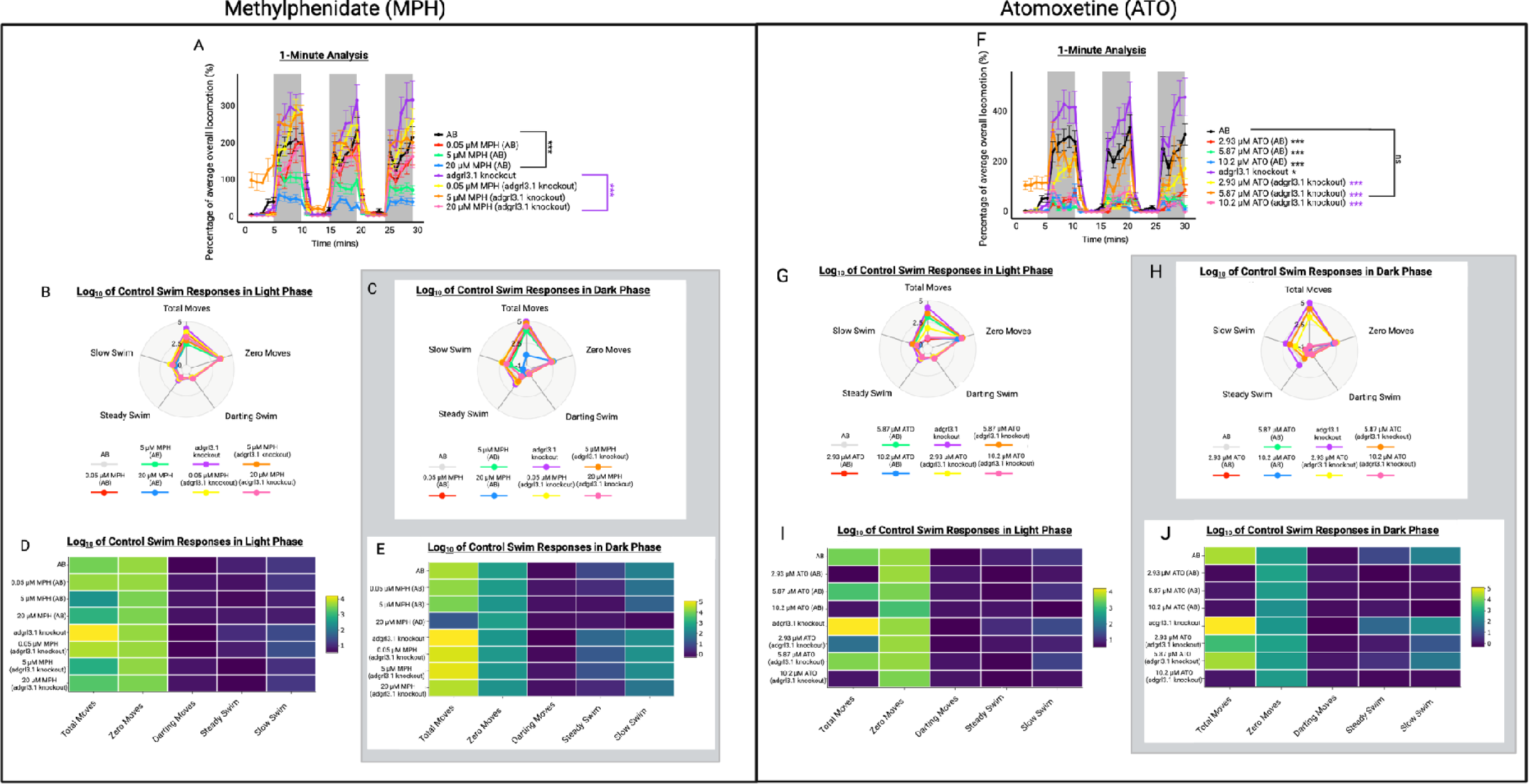
The effects of AB wild type and transgenic adgrl3.1^-/-^ larval phenotypes on light/dark locomotion as well as the potential for assessing herapeutic agents. The minute analyses (A and F) are represented as mean percentage of baseline movement over the 30-minute recording period. The heat maps (B and G) and radar charts (D and I) show light phase responses and (C and H) and (E and J) show dark phase responses.

The *adgrl3.1*^-/-^ larvae express phenotypic hyperactivity with increased dark phase locomotion compared to WT (dark phase linear mixed model analysis (strain concentration): *β* = 79.68 (SE = 31.55, *t* = 2.53) (Fig. 10). The LRT showed a significant difference between the fitted model and the null model, X2 (7) = 61.64, P < 0.0001. 5 μM MPH exposed *adgrl3.1* ^-/-^ larvae display similar movement responses to that of WT fish, with reduced total movements compared to *adgrl3.1*^-/-^ larvae. 20 μM MPH exposed *adgrl3.1*^-/-^ larvae display similar behavioural responses to 5 μM MPH exposed; however, results in significant reductions in WT total movements. These findings suggest that 5 μM MPH is optimal for reversing the hyperactive phenotype of *adgrl3.1*^-/-^ larvae without impacting WT responses compared to the other tested concentrations. These observations highlight the value of including these additional analyses and interpretations of the larval light/dark data.

Like MPH, ATO is a noradrenergic reuptake inhibitor which reduces hyperactivity in children with ADHD and has previously been shown to reverse the hyperactive phenotype of *adgrl3.1*^-/-^ larvae (Sveinsdóttir et al., 2023). However, studies have reported that a major side-effect of ATO in patients is sedation (Bastiaens, 2007). Therefore, here we examined the effects of ATO on the light/dark assay response at concentrations previously used with *adgrl3.1*^-/-^ larvae with the objective to determine the translational relevance of our assay for evaluating complex effects such as sedation (Sveinsdóttir et al., 2023).

Our findings show that like 20 μM MPH, the WT larvae exposed to ATO experienced significant reductions in locomotion compared to untreated WT larvae (linear mixed model analysis (lights effect): *β* = 121.06 (SE = 3.52, *t* = 34.43) (Fig. 10F). The LRT showed a significant difference between the fitted model and the null model, X2(7) = 104.78, P < 0.0001. *adgrl3.1*^-/-^ larvae exposed to 2.93 μM and 10.2 μM ATO had reduced dark phase locomotion compared to untreated WT fish (Fig. 10F). However, 5.87 μM ATO exposed *adgrl3.1*^-/-^ larvae had unchanged dark phase locomotion compared to untreated WT fish (Fig. 10F).

ATO exposure has a significant effect on both light and dark phase swim phenotypes (Fig. 10G-J). The radar charts and heat maps demonstrate that the major differences between the WT and *adgrl3.1*^-/-^ larvae exposed to ATO are in the total movement responses. The WT larvae exposed to all three ATO concentrations experienced significantly reduced total movement compared to untreated WT larvae in both light and dark phases (Fig. 10G-J). Comparatively, the *adgrl3.1*^-/-^ 5.87 μM ATO treated larvae had comparable total movements compared to untreated WT larvae. Our ATO observations highlight the importance of including the additional analyses for depicting adverse effects of therapeutic substances in larval zebrafish and promotes the translational relevance of our assay.

## 4. Discussion

Here, we aimed to determine the optimal conditions for the light/dark behavioural assay in 4 dpf zebrafish larvae. Our primary objective was to develop a robust and reproducible light/dark assay for 4 dpf zebrafish larvae with an in-depth analysis plan, and to validate the assay both pharmacologically, and with genetically modified larvae. The premise of this study was for providing a standardised assay for future larval research in neuropsychopharmacology and biological psychiatry.

There were several key findings here that should be highlighted. First, high levels of between-batch and inter-individual variability were found, which were corrected with normalisation. Second, we identified that a period of five-minutes for each dark and light period was optimal in terms of locomotor activity in 4 dpf larvae. A 30-minute light phase acclimation period resulted in significantly increased dark phase larval locomotion. We confirmed the pharmacological predictivity of the assay using ethanol, which caused low concentration hyperlocomotion and high concentration hypolocomotion, as predicted. Finally, the behavioural phenotypes of transgenic (adgrl3.1^-/-^) larvae were reported on, where hyperlocomotion behaviour was rescued with both MPH and ATO. We also demonstrate the translational relevance of our assay for observing potential adverse effects of therapeutic substances, with ATO exposure resulting in hypolocomotion in WT larvae. As well as validation of our assay with transgenic larvae and ethanol, we have provided a comprehensive overview of alternative methods of analysis and interpretation of the data.

In previous work, we had noted variations in larval responses between different laboratories -- for example, if larvae are transported to different sites, or when working with collaborators – even when running identical light/dark assay protocols with fish bred at the same facility (see SD 2). Larval zebrafish variability in behavioural responses is well established (Jacobs & Ryu, 2023) with both genetic and environmental factors impacting batch-batch variability (Xie et al., 2019). For instance, hatching time has a significant effect upon dark phase responsiveness in the light/dark assay; our time-of-day analysis identified no significant effects upon the behavioural responses (Rock et al., 2022). However, we did observe large variations in behavioural responses during our time-of-day analysis, which was rectified with normalisation (see SD 3). Therefore, normalisation of the data is essential (Xie et al., 2019). We established a normalisation technique where the dataset of interest is analysed as a percentage against the average baseline locomotor responses, which was able to overcome the batch variability, time-of-day variability and variability between laboratories (see SD 2). We encourage normalisation of the light/dark data for facilitating statistical comparisons and collaboration between research laboratories (Xie et al., 2019).

Development of the assay involved confirmation of the optimal transition period length and number of repetitions. Researchers have previously used a wide range of different transition periods with further variation in the number of repetitions in the assay (Table 1). Here, we evaluated the protocols employed by other researchers (Acevedo-Canabal et al., 2019; Bachour et al., 2020; Haigis et al., 2022), as well as assessing our own developed protocol. Light to dark transitions were used, as our preliminary findings showed no significant difference between light to dark or dark to light transitions for larval locomotion (SD 4). We report that the ten-minute and five-minute protocols are both suitable for assessing behavioural responses in 4 dpf zebrafish larvae. We also suggest that the five-minute protocol is superior due to its time-effectiveness, enhanced throughput and sustained increased dark phase locomotion compared to the ten-minute protocol. The larvae exposed to the two-minute transition and fifteen-minute transition protocols had reduced dark phase locomotor responses by the final dark phases compared to the first (Fig. 3B and H). This indicates that the longer paradigms are unsuitable for 4 dpf light/dark assay studies due to reduced locomotion, which hinders the potential use of the assay for assessing drug effects for example. These findings are supported by Maeda (2021) who demonstrated that larvae exposed to 15-minute intervals had gradual decreases in dark phase locomotion. They did not however observe this with the five-minute protocol of the same length. A possible reason for the reduced locomotion observed could be habituation to the dark (Tuz-Sasik et al., 2022). Habituation is non-associative learning defined here as a decrease in dark phase locomotion following repeated exposure to the light/dark transitions (Tuz-Sasik et al., 2022). In the wild, repeated exposure to harmless stimuli causes high energy expenditure and motor fatigue. Therefore, habituation ensures the fish conserves energy for survival (Tuz-Sasik et al., 2022). Another possible reason for the reduced dark phase locomotion may be exhaustion brought-on by excessive energy use (Tuz-Sasik et al., 2022). However, confirmation of these hypotheses requires further investigation.

Previous research has highlighted the importance of the length and lighting conditions of the acclimation period used prior to behavioural evaluation with fish (Makaras et al., 2021). In these studies, the effect of changing the acclimation period conditions on locomotor responses was also observed. Using our reported protocol, we identified a significant increase in dark phase locomotion with the 30-minute acclimation period (Fig. 5) as well as increased overall locomotion with light acclimation compared to dark (Fig. 6). Our evidence therefore supports the use of a 30-minute light phase acclimation period for increased locomotor responses in 4 dpf larvae.

It has been suggested that the zebrafish circadian rhythm has no influence upon light/dark assay locomotion (Krylov et al., 2021). Our findings in SD 3 confirms this with no difference in dark phase locomotion regardless of the time of day of recording. However, observations by Fitzgerald (2019), Kristofco (2016) and MacPhail (2009), disagreed with our findings. The discrepancy between observations could be methodology-dependent, which highlights the importance of the experimental parameters of the light/dark assay with zebrafish larvae (Fitzgerald et al., 2019). Slight differences in dark phase locomotion at different times of the day was observed when using the distance travelled data (SD 2A). However, this variation is removed through normalisation to baseline (SD 2B). Therefore, we strongly encourage incorporation of normalisation to baseline findings for consistent and comparable results throughout the day, alleviating the need to limit behavioural work to a certain time of day.

We aimed to further validate the assay by administering ethanol, which we selected due to its robust concentration-dependent behavioural effects (low concentration hyperactivity and high concentration hypoactivity in the dark phase (Cohen et al., 1997)), its widespread availability and common global abuse (McLellan, 2017) and its frequent use in larval light/dark assay research (Davis et al., 2021; Irons et al., 2010; MacPhail et al., 2009). The diverse and well understood neuropharmacology of ethanol (low concentration hyperactivity and high concentration hypoactivity) allowed for suitable assay validation. Low concentration ethanol hyperactivity has previously been reported in older larvae using the light/dark assay (MacPhail et al., 2009) and a similar finding was reported here with 1.0 % ethanol exposure (Fig. 7). This low concentration hyperactivity is a result of elevated dopamine levels and is commonly reported as the positive reinforcing effects of ethanol in rodents (Cohen et al., 1997). Mid-range concentrations are known to result in behavioural disinhibition in humans (Harrison et al., 2017) and similar dissociative-like swim behavioural responses have previously been reported (MacPhail et al., 2009). This dissociative-like response can be seen here in larvae exposed to 2.0 % ethanol (Fig. 7). High concentration hypoactivity stems from ethanol’s sedative-like properties in rodents and mammals, which are attributed to its GABAergic effects (Clayman & Connaughton, 2022). Like previous publications, (Irons et al., 2010; MacPhail et al., 2009), we report ethanol-induced sedative-like responses with exposure to high concentrations (Fig. 7). These observations with ethanol in the light/dark assay indicate consistent neuropharmacological mechanisms between larval zebrafish and previous findings and suggests that the assay is suitable as a first-tier model for predicting drug and chemical effects.

We further validated the assay for neuropsychopharmacology evaluation using an established transgenic line, adgrl3.1^-/-^, which was employed because previous studies had shown robust hyperactivity compared to WT (Sveinsdóttir et al., 2023). This hyperactivity was successfully reduced with exposure to both ATO and MPH (Fig. 10), which has previously been reported in ATO treated adult adgrl3.1^-/-^ zebrafish (Fontana et al., 2023) and in 6 dpf larval adgrl3.1^-/-^zebrafish treated with ATO and MPH (Lange et al., 2012). Here, we demonstrated the ability of our assay to distinctly establish more specific drug effects by demonstrating the common adverse effect, sedation, in the WT ATO exposed larvae (Bastiaens, 2007) (Fig.10). In contrast, MPH only reduced movement in WT larvae at the top concentration, consistent with its stimulant mechanism of action (Challman & Lipsky, 2000). These observations indicate this light/dark assay is appropriate for differentiating and highlighting intrinsic and potentially complex effects of compounds.

There are several ways that the larval light/dark assay data can be presented, and typically this takes the form of mean distance travelled (Haigis et al., 2022). Although this has advantages (e.g., it is quick and uses the raw data), we believe this approach has its limitations. First, as highlighted, when analysing the distance travelled data at face value there was major variation between different laboratories, and even batches within the same laboratory, using fish bred in the same facility and with identical experimental parameters. We postulate that this is associated with environmental stressors, transport, or other generic inter-batch variability (Xie et al., 2019). Secondly, different common swim phenotypes such as the freezing response cannot be accurately identified with just distance travelled data (Kalueff et al., 2013). These limitations prompted us to adopt a broader analytical approach (from Steele (2018)) and analyse responses as different swim parameters. Haigis (2022) also analysed the data in this format to produce a radar chart; however, they chose to normalise the data to control response and perform a log_2_ transformation. We similarly normalised the data to baseline responses, followed by analysis of the different swim parameters. Next, we normalised the individual swim responses to control larval response and performed a logarithmic transformation to overcome any exponential increases in swimming behaviour which was identified as impacting the readability of the radar charts and heat maps.

The swim parameters analysed here include total movement, darting movement, steady swimming movement, slow swimming movement and zero movements. We believe these swim phenotypes will facilitate more complex behavioural analysis with 4 dpf zebrafish larvae. We demonstrate that the radar charts and heat maps can distinguish more precise movement patterns and drug/genomic induced behavioural responses which cannot be ascertained solely from the line graph or bar chart (Fig. 7 and 10). Darting swimming is an important swim factor to analyse because it is an indicator of the initial dark escape/light-seeking response as well as a sign of anxiety-like responses (Haigis et al., 2022). Slow swimming responses highlight more clearly any reduced swim response whilst zero movements show the innate freezing common with zebrafish as well as complete loss of movement because of sedation, paralysis or death (Kalueff et al., 2013). A steady swim response was also included, which we postulate is continuous swimming which is not commonly observed in control 4 dpf larvae. As highlighted by, Haigis (2022), including analyses such as the radar chart permits identification of behavioural responses which may go unnoticed in standard analyses. We believe our findings highlight the radar chart and heat map value in analysis and strongly encourage the inclusion of both to facilitate the readability and interpretation of future light/dark larval behavioural research.

## 5. Potential applications

Our validated assay promotes and encourages the use of 4 dpf zebrafish larvae for neuropsychopharmacology experimentation, which is beneficial from a 3R’s of animal research perspective (Sneddon et al., 2017). The model can be used for studying potential therapeutic agents. For example, we used adgrl3.1^-/-^ transgenic larvae to observe the effects of MPH and ATO, known anti-ADHD drugs, (Yang et al., 2018), as well as assessing the genetic behavioural phenotypes. There is also the potential for the model to be used for toxicology and ecotoxicology research (Haigis et al., 2022). Previous research has identified the use of the light/dark assay for assessing anxiogenic and anxiolytic substances (Facciol et al., 2019). However, here we suggest the assay can be used to screen for a range of pharmacologically induced behavioural parameters, including hyperlocomotion and sedation. Additionally, we believe a potential application for the assay would be to assess environmental effects on behaviour (e.g., water condition) (Rock et al., 2022). These potential applications we believe highlight the unique flexibility of this larval assay.

## 6. Limitations

We acknowledge several limitations. First, although we highlight excellent translational potential of the 4 dpf behavioural data, they remain a preliminary model only. Analysis of more complex behaviours is likely not possible at this age due to their underdeveloped CNS (Blader, 2000). Second, this method is reliant upon accurate behavioural tracking and therefore the tracking settings may need adjustment prior to investigation. Third, our new analytical method is more complex than simplistic averaging of distance travelled; however, the custom R script available on OSF overcomes this and facilitates analysis.

## 7. Conclusions

Here we present a standardised light/dark behavioural assay for use in 4 dpf zebrafish larvae, which was validated using exposure to ethanol with wild type AB and exposure to ADHD drugs with adgrl3.1^-/-^ transgenic larvae. We report that the five-minute transition period is optimal for consistent and reproducible locomotor responses and that an acclimation period is not required for robust responses to be seen, which improves the throughput of the assay. We present the influence of time of day on 4 dpf larval responses whilst providing a comprehensive analysis of different methods of interpretation and presentation of the behavioural data. Finally, this work has strong 3Rs applications. Despite some limitations due to their reduced ability to determine complex behaviours, we strongly encourage the incorporation of 4 dpf studies prior to adult or mammalian neuropsychopharmacology studies to reduce the overall number of protected animals used in studies.

## Abbreviations

ATO: atomoxetine
dpf: days post-fertilization
LRT: likelihood ratio test
MPH: methylphenidate
SD: supplementary data
WT: wild-type

## 8. Declarations/Acknowledgments

### Ethics approval

All experiments were approved by the University of Portsmouth Animal Welfare and Ethical Review Board, and under license from the UK Home Office (Animals [Scientific Procedures] Act, 1986, license number PP8708123).

### Consent to participate

Not applicable.

### Consent for publication

Not applicable.

### Availability of data and materials

All data and analysis materials are available on the Open Science Framework. https://osf.io/k7y2c/?view_only=61e23ea4bb2d452dab76646931e3f332

### Code availability

Not applicable.

### Authors’ contributions

Conceptualization: CH, JK, MP. Formal analysis: CH. Funding acquisition: JK, MP. Investigation: CH. Methodology: CH, JK, MP. Project administration: MP. Resources: JK, MP. Supervision: JK, MP. Visualization: CH. Writing – original draft preparation: CH. Writing – review & editing: CH, JK, MP.

## 10. Supplementary Data

SD 1: Previous studies have shown that the size of well plate used for larval behavioural research significantly impacts the behavioural response (Widrick et al., 2023). However, here we demonstrate that any well plate size is suitable for 4 dpf zebrafish larval light/dark behavioural research (SD 1). No significant difference was seen following normalisation in both light and dark phases for the well plates. However, the lack of movement in the light phase with the 96-well plate is extreme and these floor effects means it is hard to observe any movement suppression with pharmacological interventions (SD 1). Therefore, we selected the 48-well plate for further analysis due to its enhanced throughput compared to the 24-well plate and improved time-efficiency for plating compared to the 96-well plate as well as increased reliability. However, our evidence suggests that well plate size should not impact the behavioural results seen if normalisation is performed.

**SD 1:**
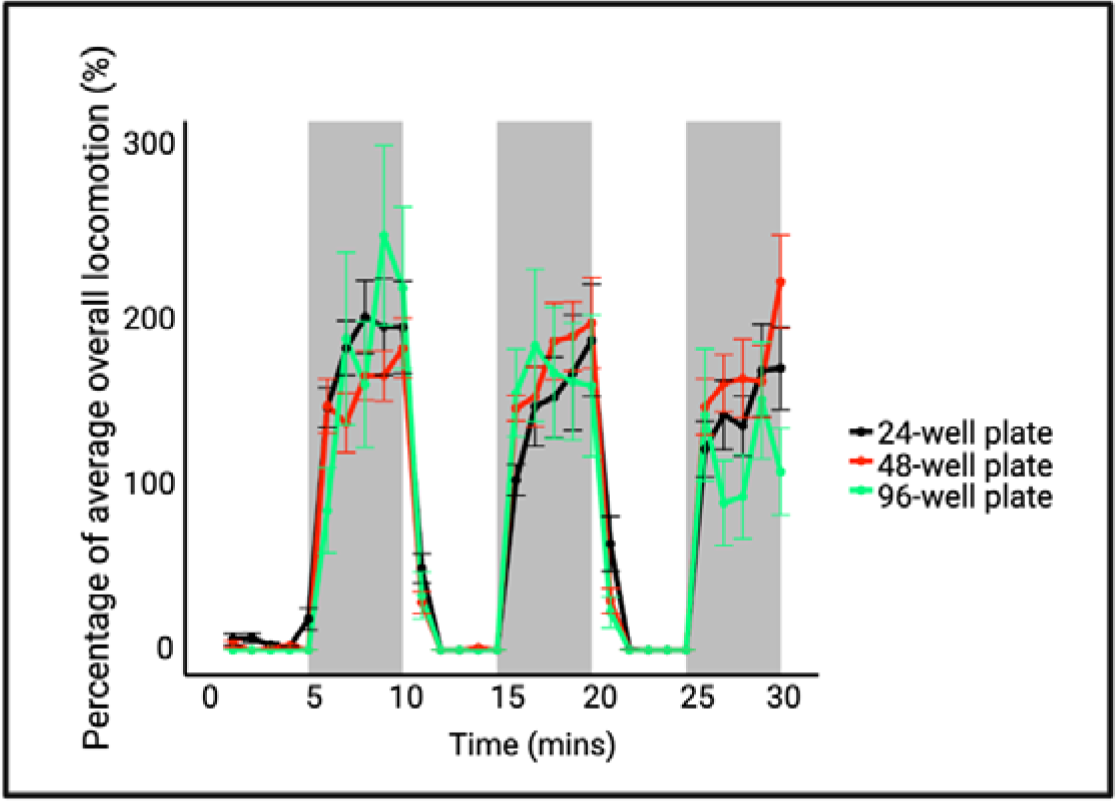
The minute analysis comparing the average locomotion for 24-well plates, 48-well plates and 96-well plates.

SD 2: We noted substantial differences between our two laboratories running identical light/dark protocols with fish obtained from the same fish facility (University of Portsmouth). We highlight the importance of normalising the data before analysis rather than using the distance travelled data (SD 2A-F). Baseline dark phase locomotion was significantly different between the two laboratories using distance travelled data (one-way ANOVA: F_(1, 30)_ = 14.3, p < 0.0001) (SD 2C). No difference was observed between the light phases (one-way ANOVA: F_(1, 30)_ = 1.028, p = 0.319). Comparatively, following normalisation to baseline no significant difference in locomotion was observed for either dark phase or light phase.

**SD 2:**
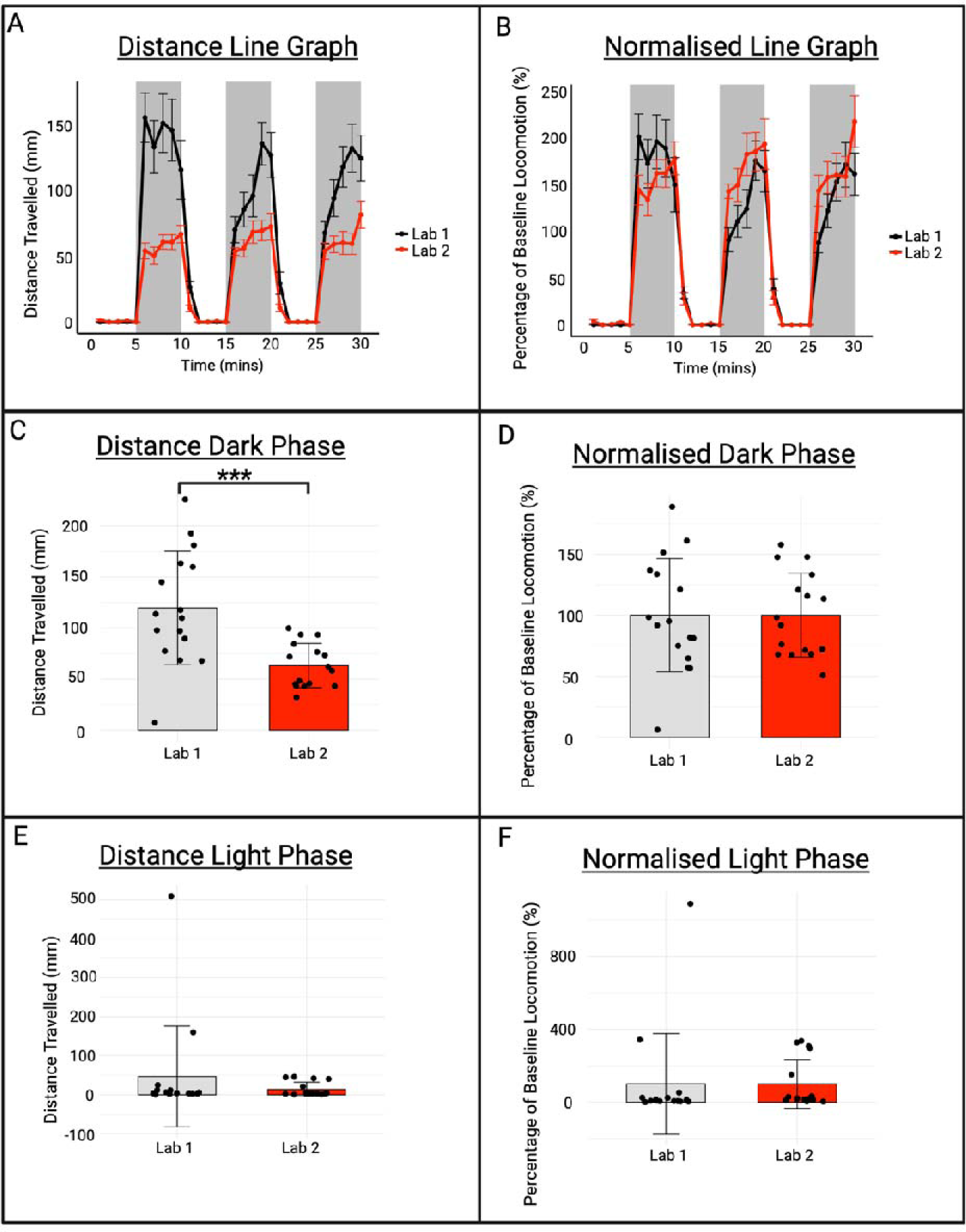
Comparison of baseline data obtained from our two separate laboratories using fish from the same fish facility. The data is presented as either distance travelled data as a line graph (A) or normalised data as a line graph (B). The dark and light phase responses for distance travelled (C and E) and normalised data (D and F) are also shown.

SD 3: It has previously been reported that the time of day affects larval zebrafish behavioural responses using the light/dark assay (Fitzgerald et al., 2019; Kristofco et al., 2016; MacPhail et al., 2009). We observed no significant effect of time of (working) day (morning (09:00 – 11:59), afternoon (12:00 and 14:29) and evening (14:30– 17:00)) on larval light/dark assay (SD3 A and B). There is slight variability in dark phase locomotion using the distance travelled data (one-way ANOVA: F_(2, 45)_ = 1.183, p = 0.316); however, this is fully removed following normalisation (one-way ANOVA: F_(2, 45)_ = 0, p = 1).

**SD3.**
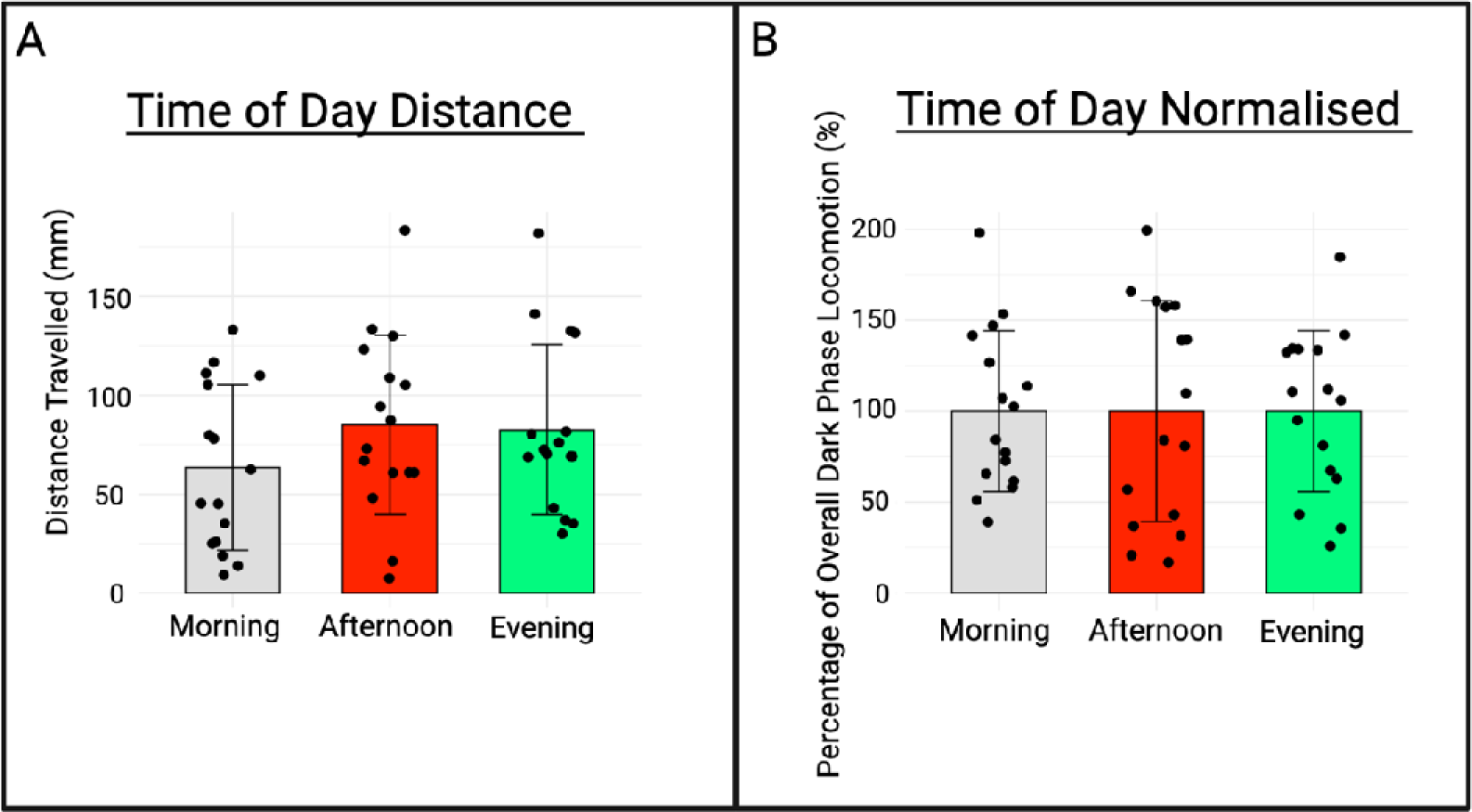
Time of day of the light/dark assay recording affects behavioural responses observed in 4 dpf zebrafish larvae represented as (A) distance travelled data and (B) normalised to baseline data.

SD4: We observed no significant difference in locomotion starting with light to dark vs dark to light transitions in the light phase or dark phase (SD 4C and D).

**SD 4:**
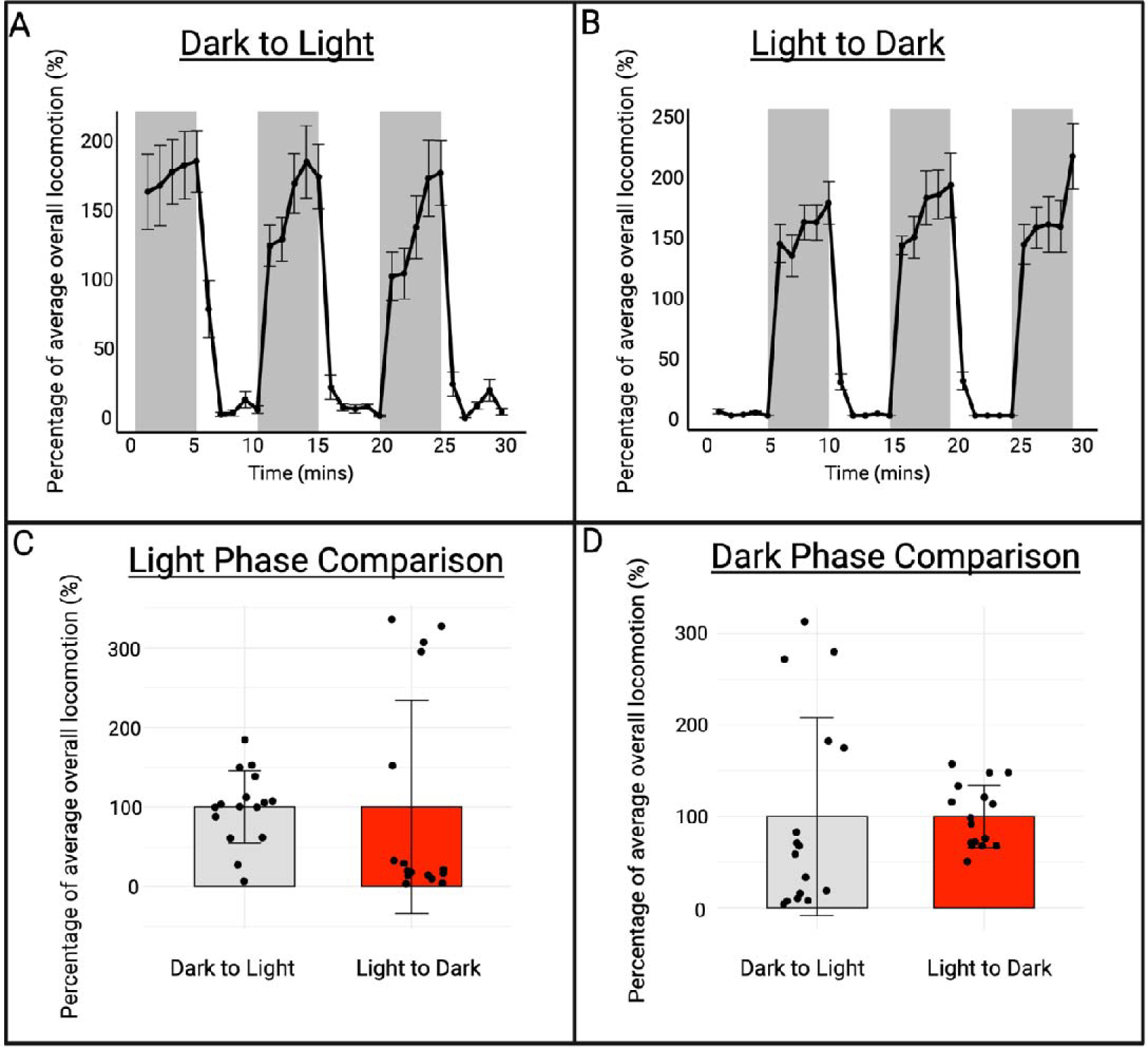
The behavioural effects of running the assay light to dark (A) or dark to light (B) reported as percentage of baseline recording in minutes. The mean light phase (C) and dark phase (D) comparison is reported.

