## Supplementary data for "A Unified Approach to Investigating 4 dpf Zebrafish Larval Behaviour through a Standardised Light/Dark Assay"

1. **Supplementary Data**

**SD 1:** Previous studies have shown that the size of well plate used for larval behavioural research significantly impacts the behavioural response (Widrick et al., 2023). However, here we demonstrate that any well plate size is suitable for 4 dpf zebrafish larval light/dark behavioural research (SD 1). No significant difference was seen following normalisation in both light and dark phases for the well plates. However, the lack of movement in the light phase with the 96-well plate is extreme and these floor effects means it is hard to observe any movement suppression with pharmacological interventions (SD 1). Therefore, we selected the 48-well plate for further analysis due to its enhanced throughput compared to the 24-well plate and improved time-efficiency for plating compared to the 96-well plate as well as increased reliability. However, our evidence suggests that well plate size should not impact the behavioural results seen if normalisation is performed.

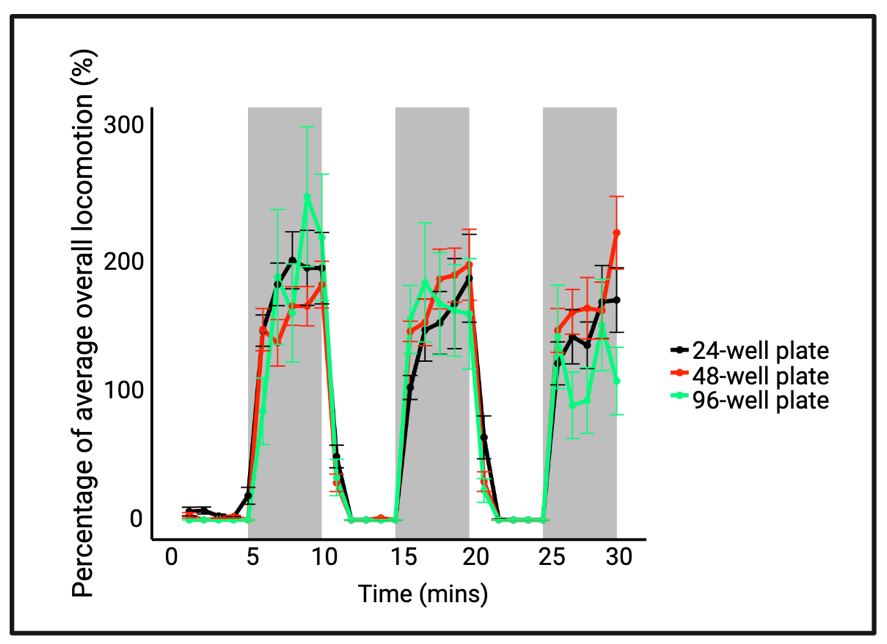

**SD 1: The minute analysis comparing the average locomotion for 24-well plates, 48-well plates and 96-well plates.**

**SD 2:** We noted substantial differences between our two laboratories running identical light/dark protocols with fish obtained from the same fish facility (University of Portsmouth). We highlight the importance of normalising the data before analysis rather than using the distance travelled data (*SD* 2A-F). Baseline dark phase locomotion was significantly different between the two laboratories using distance travelled data (one-way ANOVA: F_(1, 30)_ = 14.3, *p* < 0.0001) (*SD* 2C). No difference was observed between the light phases (one-way ANOVA: F_(1, 30)_ = 1.028, *p* = 0.319). Comparatively, following normalisation to baseline no significant difference in locomotion was observed for either dark phase or light phase.

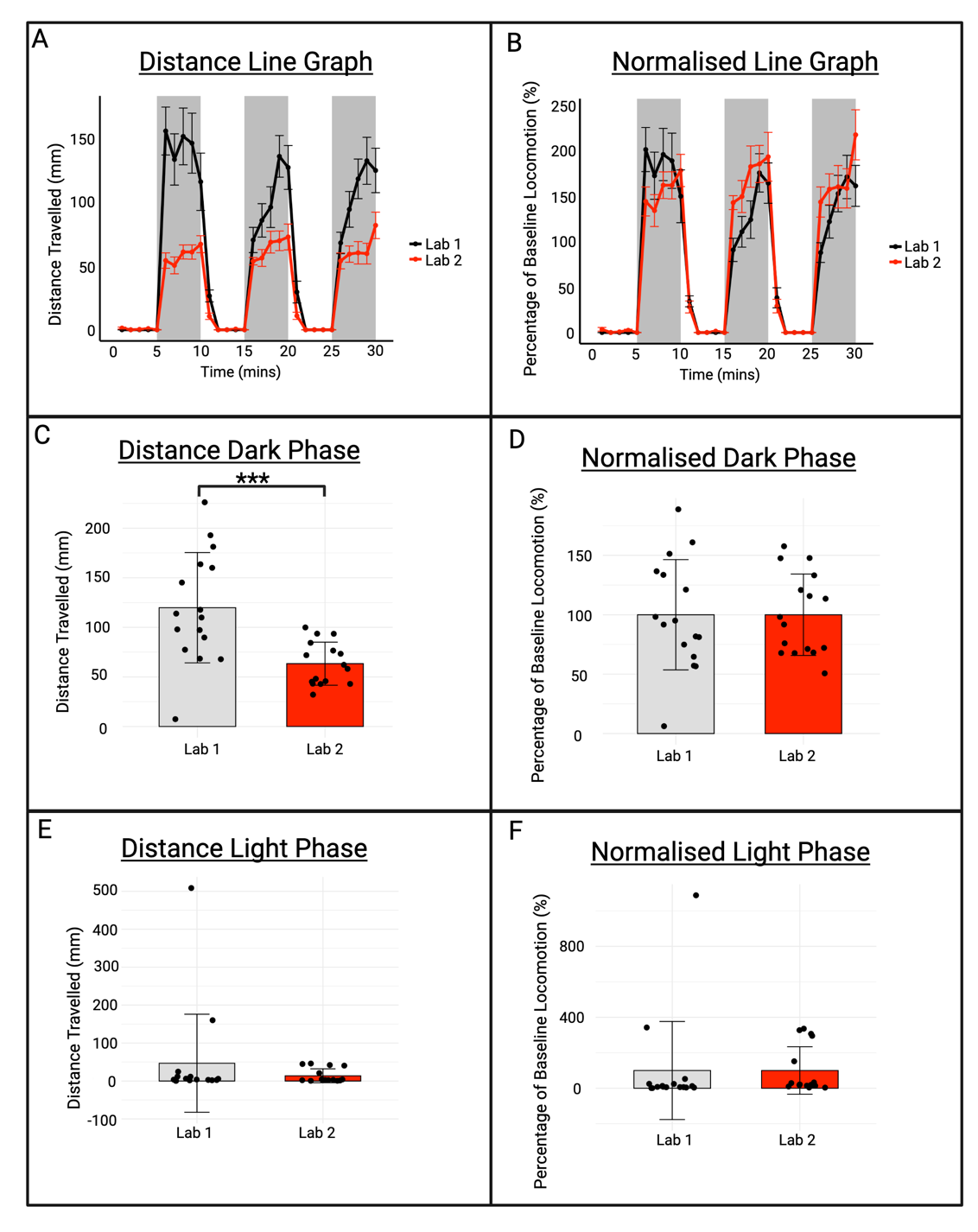

**SD 2: Comparison of baseline data obtained from our two separate laboratories using fish from the same fish facility.** The data is presented as either distance travelled data as a line graph (A) or normalised data as a line graph (B). The dark and light phase responses for distance travelled (C and E) and normalised data (D and F) are also shown.

**SD 3:** It has previously been reported that the time of day affects larval zebrafish behavioural responses using the light/dark assay (Fitzgerald et al., 2019; Kristofco et al., 2016; MacPhail et al., 2009). We observed no significant effect of time of (working) day (morning (09:00 – 11:59), afternoon (12:00 and 14:29) and evening (14:30– 17:00)) on larval light/dark assay (SD3 A and B). There is slight variability in dark phase locomotion using the distance travelled data (one-way ANOVA: F_(2, 45)_ = 1.183, *p* = 0.316); however, this is fully removed following normalisation (one-way ANOVA: F_(2, 45)_ = 0, *p* = 1).

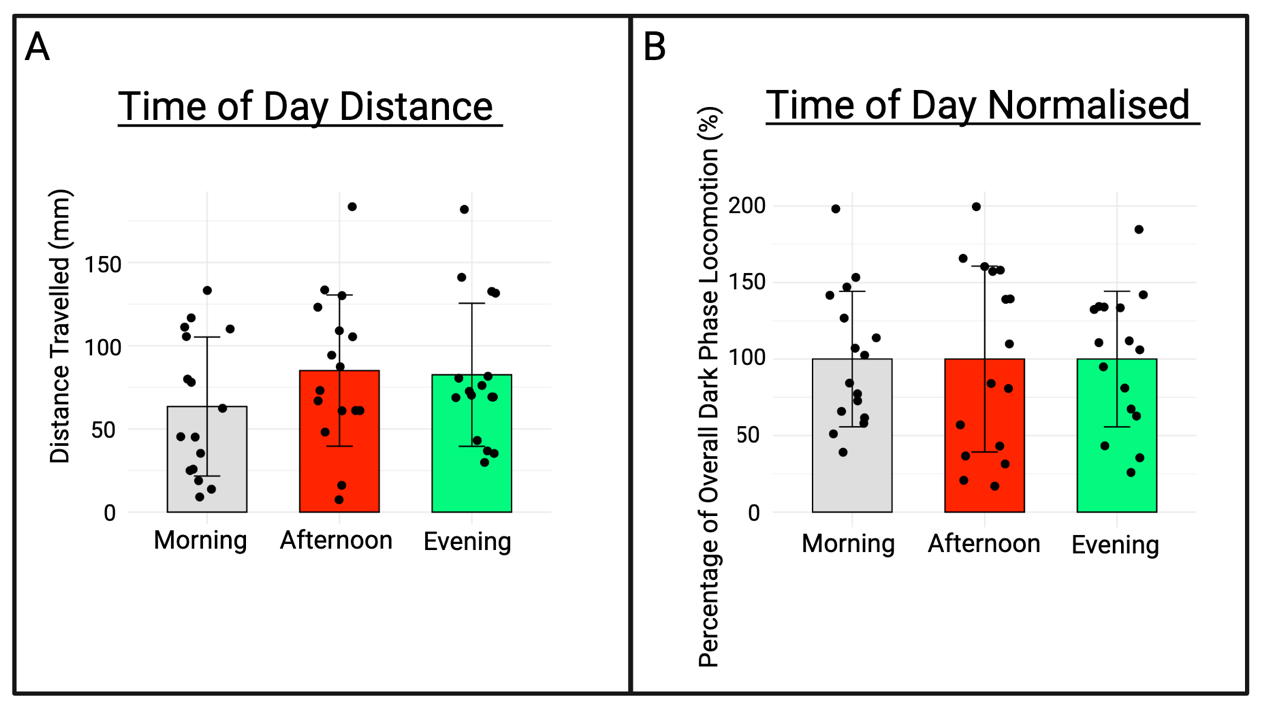

**SD3. Time of day of the light/dark assay recording affects behavioural responses observed in 4 dpf zebrafish larvae represented as (A) distance travelled data and (B) normalised to baseline data.**

**SD4:** We observed no significant difference in locomotion starting with light to dark vs dark to light transitions in the light phase or dark phase (*SD* 4C and D).

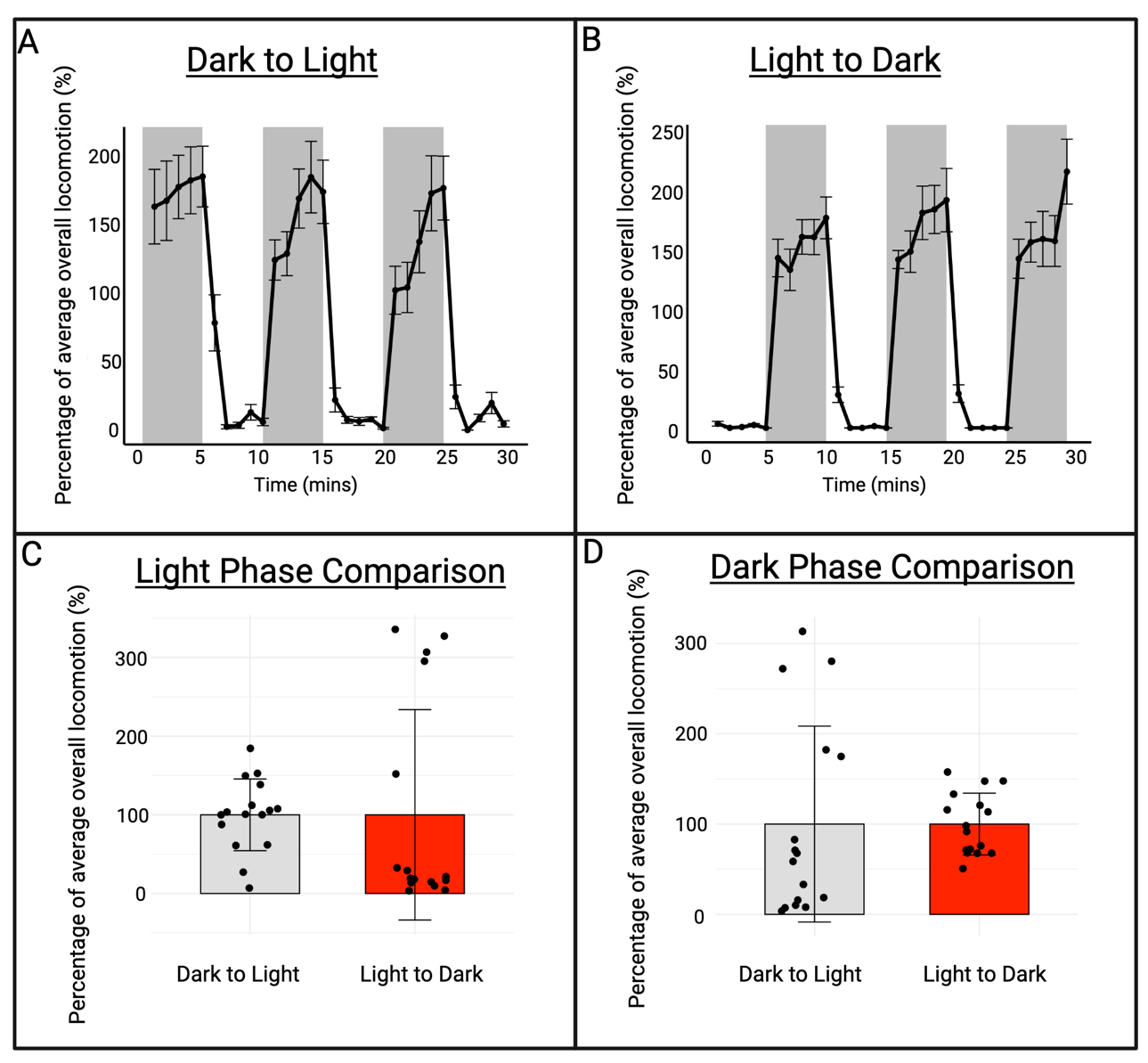

**SD 4:** The behavioural effects of running the assay light to dark (A) or dark to light (B) reported as percentage of baseline recording in minutes. The mean light phase (C) and dark phase (D) comparison is reported.
